# Periodic ER-plasma membrane junctions support long-range Ca^2+^ signal integration in dendrites

**DOI:** 10.1101/2024.05.27.596121

**Authors:** Lorena Benedetti, Ruolin Fan, Aubrey V. Weigel, Andrew S. Moore, Patrick R. Houlihan, Mark Kittisopikul, Grace Park, Alyson Petruncio, Philip M. Hubbard, Song Pang, C. Shan Xu, Harald F. Hess, Stephan Saalfeld, Vidhya Rangaraju, David E. Clapham, Pietro De Camilli, Timothy A. Ryan, Jennifer Lippincott-Schwartz

## Abstract

Neuronal dendrites must relay synaptic inputs over long distances, but the mechanisms by which activity-evoked intracellular signals propagate over macroscopic distances remain unclear. Here, we discovered a system of periodically arranged endoplasmic reticulum-plasma membrane (ER-PM) junctions tiling the plasma membrane of dendrites at ∼1 μm intervals, interlinked by a meshwork of ER tubules patterned in a ladder-like array. Populated with Junctophilin-linked plasma membrane voltage-gated Ca^2+^ channels and ER Ca^2+^-release channels (ryanodine receptors), ER-PM junctions are hubs for ER-PM crosstalk, fine-tuning of Ca^2+^ homeostasis, and local activation of the Ca^2+^/calmodulin-dependent protein kinase II. Local spine stimulation activates the Ca^2+^ modulatory machinery facilitating voltage-independent signal transmission and ryanodine receptor-dependent Ca^2+^ release at ER-PM junctions over 20 μm away. Thus, interconnected ER-PM junctions support signal propagation and Ca^2+^ release from the spine-adjacent ER. The capacity of this subcellular architecture to modify both local and distant membrane-proximal biochemistry potentially contributes to dendritic computations.

**Highlights:**

- Periodic ER-PM junctions tile neuronal dendritic plasma membrane in rodent and fly.
- ER-PM junctions are populated by ER tethering and Ca^2+^ release and influx machinery.
- ER-PM junctions act as sites for local activation of CaMKII.
- Local spine activation drives Ca^2+^ release from RyRs at ER-PM junctions over 20 μm.

## Introduction

Neuronal dendrites execute computations in the brain using repertoires of voltage- and ligand-gated ion channels to locally influence the output of different synaptic inputs across the plasma membrane (PM)^1^. Extensive research has elucidated the mechanisms of electrical conduction along the plasma membrane of dendrites, but how spatially and temporally confined intracellular messengers produced by these channels, such as Ca^2+^, which is dissipated or buffered over the nanoscale^2–5^, affects membrane-proximal biochemistry is unclear. The answer to this question is crucial for understanding how information from distinct synaptic inputs at dendritic spines is summated and processed in the brain.

The endoplasmic reticulum (ER) is the cell’s largest organelle, comprising a surface area many times that of the plasma membrane. Extending from the perinuclear area to the axon, presynaptic terminals, dendrites, and dendritic spines in neurons^6^, the ER has been proposed to be central for orchestrating the integration and transmission of intracellular signaling within neurons^7^. Indeed, mutations in intrinsic ER proteins are associated with various neuropathologies, including cognitive dysfunction, familial Alzheimer’s disease, epilepsy, hereditary spastic paraplegia, and amyotrophic lateral sclerosis^8–14^, with neuronal pathology often manifesting as alterations in ER structure^15,16^. In dendrites, the ER has specialized characteristics likely to be crucial for proper neuronal function, including Ca^2+^ release channels that can increase or reduce synaptic efficacy distant from the stimulation site^7,17–21^. In addition, ER contact sites with the PM^22,23^ have been implicated as intracellular signaling hubs in neuronal cell bodies^24–31^. The ER is an excellent candidate for intracellular Ca^2+^ signaling propagation in dendrites; however, its structure and organization are still poorly understood. This is due to several factors, including the ER’s extreme dynamism in cells^32^; the ER’s narrow caliber and tremendous length in neuronal processes^33,34^; limited availability of electron microscopy (EM) reconstructions of ER across extended regions of neurons^23,35–37;^ and incomplete information on the subcellular distribution of key ER proteins involved in Ca^2+^ signaling.

Here, we resolve dendritic ER’s structure/function in relation to neuronal signaling using a combination of multi-color super-resolution microscopy, focused ion beam scanning electron microscopy (FIB-SEM), and biochemical/physiological methods. We show that ER and PM proteins involved in Ca^2+^ signaling are concentrated at ER-PM junctions arranged in a periodic pattern along the dendrite. The ER-PM junctions act as hubs of Ca^2+^ influx and release machineries, whose composition is modulated by neuronal activity. Ca^2+^ influx and release at ER-PM junctions, in turn, support local activation of Ca^2+^/calmodulin-dependent kinase II (CaMKII). Using endogenous tagging approaches, we found that Junctophilin-3 (JPH3), whose muscle isoforms (e.g., JPH1,2) help build sarcoplasmic reticulum-PM contact sites^38–40^, populates dendritic ER-PM contact sites. The molecular partnership between JPH3 and Ca^2+^ influx and release machineries is key to this subcellular organization, as shRNA-mediated knockdown of junctophilins eliminated voltage-gated calcium channel (Ca_V_) recruitment to ER-PM junctions. Focal glutamate uncaging experiments coupled with ER Ca^2+^ imaging further revealed that local spine stimulation could elicit RyR-dependent Ca^2+^ release from the ER over length scales that span more than 20 μm.

The pattern of ER Ca^2+^ release sites coincided with the spacing of ER-PM junction intervals. Together, the results reveal a system of periodically arranged ER-PM contact sites in dendrites, conserved both in vertebrate and invertebrate neurons *in vivo*, that allows individual synaptic inputs to influence membrane-proximal biochemistry at distances that greatly exceed the dimensions of individual spines, potentially allowing long-distance crosstalk between synaptic inputs.

## Results

### A periodic array of ER-PM junctions interconnects ER and PM of dendrites both *in vitro* and *in vivo* in vertebrate and invertebrate neurons

We used 3D lattice-SIM microscopy to explore dendritic ER organization in rat hippocampal neuronal cultures co-expressing the ER marker HaloTag-Sec61β or mEmerald-Sec61β and the PM marker mScarlet-CAAX2. Examination of the entire dendrite (Figure 1A) revealed a ladder-like ER morphology in proximal and medial dendrites (Figure 1B), with each rung of the ER ladder ending at sites close to the PM (Figure 1C, proximal and medial), and a simpler ER morphology in distal dendrites comprised of a single ER tubule (Figure 1C, distal). The ladder-like ER arrays in proximal and medial dendrites were comprised of longitudinal tubules interconnecting short transversal tubular elements that intersected the PM at perpendicular angles. Proximal dendrites, with wider diameters, had more longitudinal and interconnecting transversal ER elements than medial dendrites.

**Figure 1:**
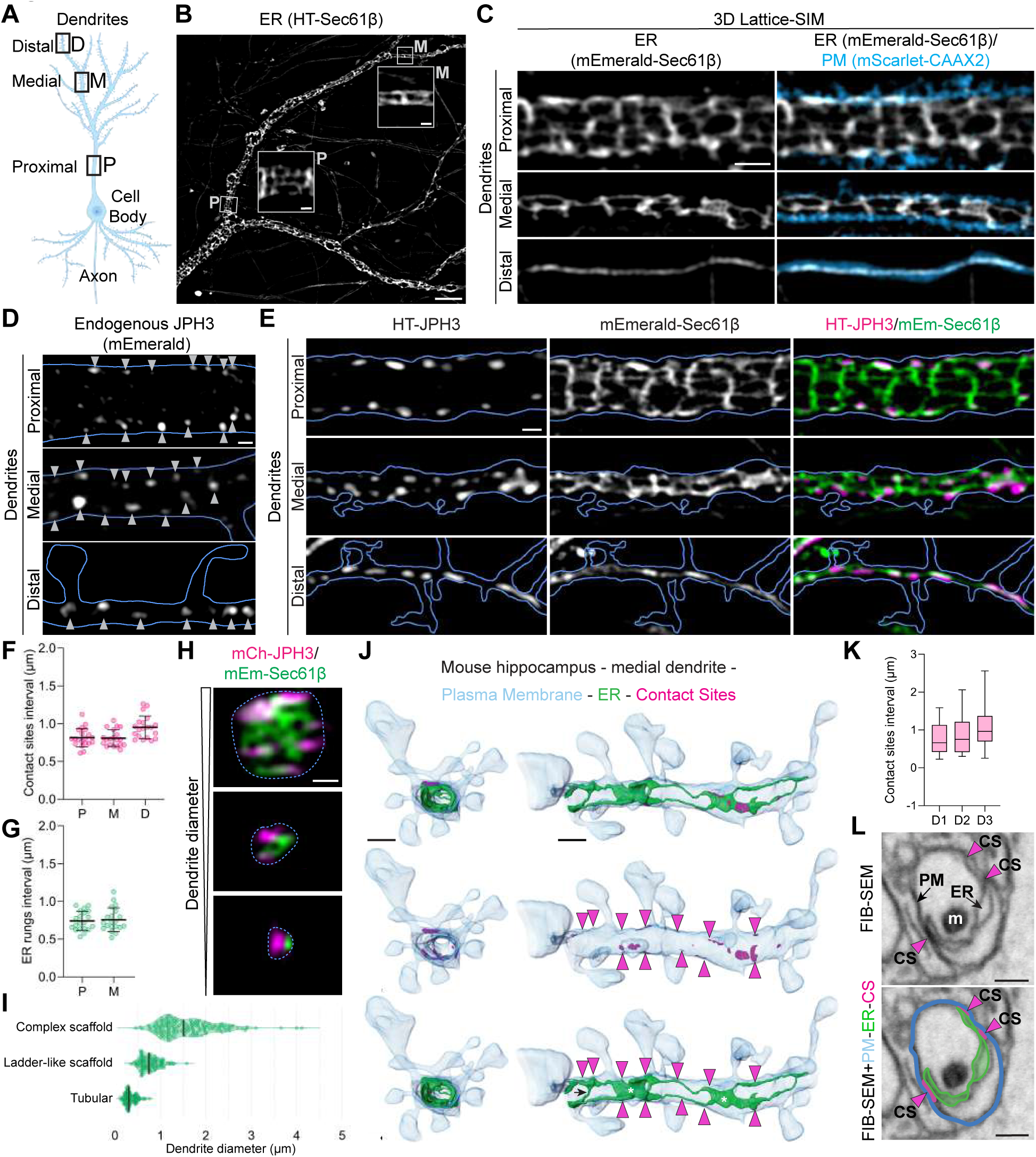
ER – PM contact sites tile the plasma membrane of hippocampal neurons at regular intervals *in vitro* and *in vivo*. A) Schematic illustration of a primary pyramidal hippocampal neuron emphasizing distinct dendritic regions: proximal dendrites (0.5 to 4.1 μm in diameter), medial dendrites (0.3 to 1.2 μm in diameter), and distal dendrites (0.1 to 0.76 μm in diameter). B) Lattice-SIM image portrays the organization of dendritic ER (expression of HaloTag-Sec61β labeled with JF635 HaloTag-ligand) in primary rat hippocampal neuron at 21 DIV. In proximal (P) and medial (M) dendritic segments, we observed a ladder-like organization of the ER, as highlighted in the corresponding insets. Scale bar: 5 μm main panel, 0.5 μm insets. C) Lattice-SIM images of proximal, medial and distal neuronal dendrites expressing mEmerald-Sec61β (ER; grayscale) and mScarlet-CAAX2 (PM; blue). Scale bar: 1 μm. D) Endogenous distribution of JPH3 (gene targeted insertion of mEmerald) in proximal, medial and distal dendrites. The blue outline highlights the outer PM signal (mTagBFP2-CAAX2). Gray arrowheads indicate the position of endogenous JPH3 clusters. Scale bar: 1 μm. E) Lattice-SIM images of proximal, medial and distal neuronal dendrites expressing HaloTag-Junctophilin-3 (HT-JPH3, labeled with JF635 HaloTag-ligand), mEmerald-Sec61β, mScarlet-CAAX2. The blue outline in each panel represents the outer PM signal (mScarlet-CAAX2). Scale bar: 1 μm. F) Average HT-JPH3 contact sites interval along the dendritic plasma membrane in proximal (0.82 ± 0.12 μm), medial (0.81 ± 0.11 μm) and distal (0.95 ± 0.15 μm) dendritic regions (mean ± SD, n = 20 neurons, N = 3 animals). G) Average interval between adjacent transversal ER tubular connections in proximal (0.74 ± 0.13 μm) and medial (0.76 ± 0.16 μm) dendrites (mean ± SD, n = 20 neurons, N = 3 animals). H) Orthogonal views of complete dendritic cross-sections captured with high-resolution Airyscan z-stacks. The dotted line delineates the process boundary. Scale bar: 500 nm. I) Plot illustrating the relationship between ER organization complexity and dendritic process diameter. In proximal and larger medial dendrites (>1.61 ± 0.63 μm in diameter, n = 165, N = 20 cells) the ER is organized as a complex scaffold. In medial dendrites, ranging between 0.35 and 1.58 μm (n = 113, N = 20 cells) in diameter, the ER is organized as a simple ladder-like scaffold. In distal dendrites, with a diameter of approximately 0.34 ± 0.12 μm, the ER appears as a single tubule (n = 113, N = 20 cells) (N= 3 animals). J) Orthogonal and longitudinal views of ER (green), plasma membrane (blue) and contact sites (magenta) segmented from FIB-SEM datasets of mouse hippocampus medial dendrites. Magenta arrowheads indicate contact sites anchoring ER tubular connections (black arrow). Asterisks indicate flattened ER cisternae. Scale bar: 1 μm. K) Box-plot of contact sites interval measured in 3 FIB-SEM segmentations of mouse hippocampus medial dendrites. (Dendrite 1 (D1) = 0.78 ± 0.43 μm, n = 16; Dendrite 2 (D2) = 0.87 ± 0.53 μm, n= 11; Dendrite 3 (D3) = 1.07 ± 0.60 μm n = 22; mean ± SD). L) FIB-SEM orthoslice (top panel) and the segmentation of endoplasmic reticulum (ER, green), plasma membrane (PM, blue) and contact sites (CS, magenta arrowheads) (bottom panel) of mouse hippocampus medial dendrite showing ER tubules anchored to opposing sides of the dendritic PM. Junctional sites contain electron-dense material. (m, mitochondrion). Scale bar: 500 nm.

Since the rungs of the laddered ER arrays anchored next to the PM (Figure 1C), we tested whether these sites contained ER-PM junctional proteins, marking them as ER-PM contact sites. One neuron-specific junctional protein is JPH3^41^, an ER-resident member of the Junctophilin family of proteins known to establish and maintain ER-PM junctions (i.e., ER-PM contact sites)^42^. We labeled JPH3 with the fluorescent protein mEmerald at its endogenous locus in primary rat hippocampal neurons^43^ (see Materials and Methods and Figure S1A-F for a detailed description of the knock-in strategy and validation). Examining neurons co-expressing JPH3 mEmerald and the PM marker mTagBFP2-CAAX2 revealed puncta of JPH3 close to the PM in both cell bodies (Figure S2A, Movie 1) and dendrites (Figure 1D). JPH3 puncta juxtaposed to the PM were widely localized along the entire dendritic arbor (Figure 1D) and were distributed at evenly spaced intervals of ∼1 μm (0.9 ± 0.1 μm; Figure S2B-D and Figure S1H-K) across all dendritic regions examined (see Supplementary Materials for discussion of spacing quantification).

To understand JPH3 puncta distribution in relation to ER and PM organization, we examined entire dendrites by 3D lattice-SIM microscopy in neurons co-expressing HaloTag-JPH3 (HT-JPH3) a fusion construct capable of recapitulating endogenous JPH3 distribution (Figure S2E), with mEmerald-Sec61β to label the ER and mScarlet-CAAX2 to label the PM. We found HT-JPH3 puncta were localized at sites of ER-PM contacts, specifically at the ends of each rung of the ladder-like ER array (Figure 1E). As found for endogenously expressed JPH3, HT-JPH3 puncta were consistently spaced at ∼0.8 ± 0.1 μm intervals along the entire dendritic arbor, including proximal, medial, and distal dendritic segments (Figure 1F, Figure S2F) and among all analyzed neurons (Figure S2G and Figure S2L-O). Intervals between ER rungs matched well with the distances between HT-JPH3 puncta (Figure 1G). A comprehensive analysis of the positioning of JPH3 contacts relative to dendritic spines revealed that when the ER protrudes into the spine head, a contact site is positioned within 600 nm of the spine neck. When the ER is absent from the spine, a JPH3-coordinated junction is found within 1 μm (Figure S2P-Q). Thus, the molecular machinery localized at these contacts might be affected by spine stimulation.

Imaging HT-JPH3 contact sites in relation to the ER in orthogonal views, we found that the ER was comprised of a mesh of anastomosing tubules attached to multiple JPH3-contact sites distributed in a ring-like fashion around the PM (Figure 1H). This observation suggested an ER anchoring role of JPH3-contact sites. Notably, narrowing of the dendritic diameter led to a reduction in the complexity of ER organization (Figure 1I) concomitant with fewer JPH3-contact sites securing the ER to the PM (Figure 1H). This occurred without any change in the spacing of contact sites longitudinally along the PM (Figure 1E and Figure 1F), hinting that such spacing was highly conserved.

We next tested whether the arrangements of ER and ER-PM contact sites seen *in vitro* could be observed *in vivo* by performing FIB-SEM and ER/PM segmentation in dendritic segments from neurons of the mouse hippocampus (Figure 1J-L). Examining ER, PM, and ER-PM contact sites in a FIB-SEM image of a medial dendrite (identified by the presence of spines) revealed a regular pattern of ER-PM contact sites along the dendrite (see magenta arrowheads). Quantification from 3-D segmentations of three different dendrites showed that the average distance between consecutive ER-PM contacts using ER tubules as a guide was ∼1 μm (0.9 ± 0.1 μm; Figure 1K), resembling ER-PM contact site intervals seen in dendrites from rat hippocampal neuronal cultures. The ER in these cells extended as a continuous network of narrow tubules beneath the PM, often connected by short tubular links (as shown by a black arrow), with small ER tubules and flattened ER cisternae (indicated by asterisks) contacting the PM at regular intervals (Figure 1J). A cross-sectional view from a FIB-SEM orthoslice showed opposing contact sites on either side of the dendritic shaft connected by an ER tubule (Figure 1L).

We also examined ER and ER-PM contact site organization in FIB-SEM datasets from invertebrate neurons, specifically the *Drosophila melanogaster* central nervous system^44^, focusing on the mushroom body (MB), a key area for associative learning in the fly brain^45,46^ (Figure S3A). As found in dendrites from mouse brain and cultured rat neurons, the ER in fly neurites showed a regular, laddered organization, with each rung ending at sites close to the PM (Figure 2A-D, Movie 2). The rungs of the ER ladders in large fly neurites were more elaborate compared with ER ladder morphology seen in thin medial mouse and rat hippocampal dendrites, consisting of ER fenestrae and anastomosing ER tubules (Figure 2A and Figure 2B) reminiscent of the appearance of the ER rungs in rat proximal dendrites (Figure 1H, upper panel). Like the rungs of ER ladders seen in hippocampal dendrites, the rungs of ER ladders in fly neurites were distributed at regular intervals (∼1.5 µm), as seen in longitudinal views (Figure 2A) and in 3D-segmentations (Figure 2B). They also contacted the PM at perpendicular angles (Figure 2C and Figure 2D). Close examination of these contact sites showed electron-dense material at the sites of tight interaction (within <30 nm) (Figure 2C). In addition to regularly spaced ER-PM contacts (indicated by magenta arrowheads and boxes), we observed larger, non-rung associated contact sites in fly neurites (Figure 2D, asterisks), where they formed from ER tubules running longitudinally beneath the PM, as was also seen in rat and mouse hippocampal dendrites (Figure S3B, asterisks).

**Figure 2:**
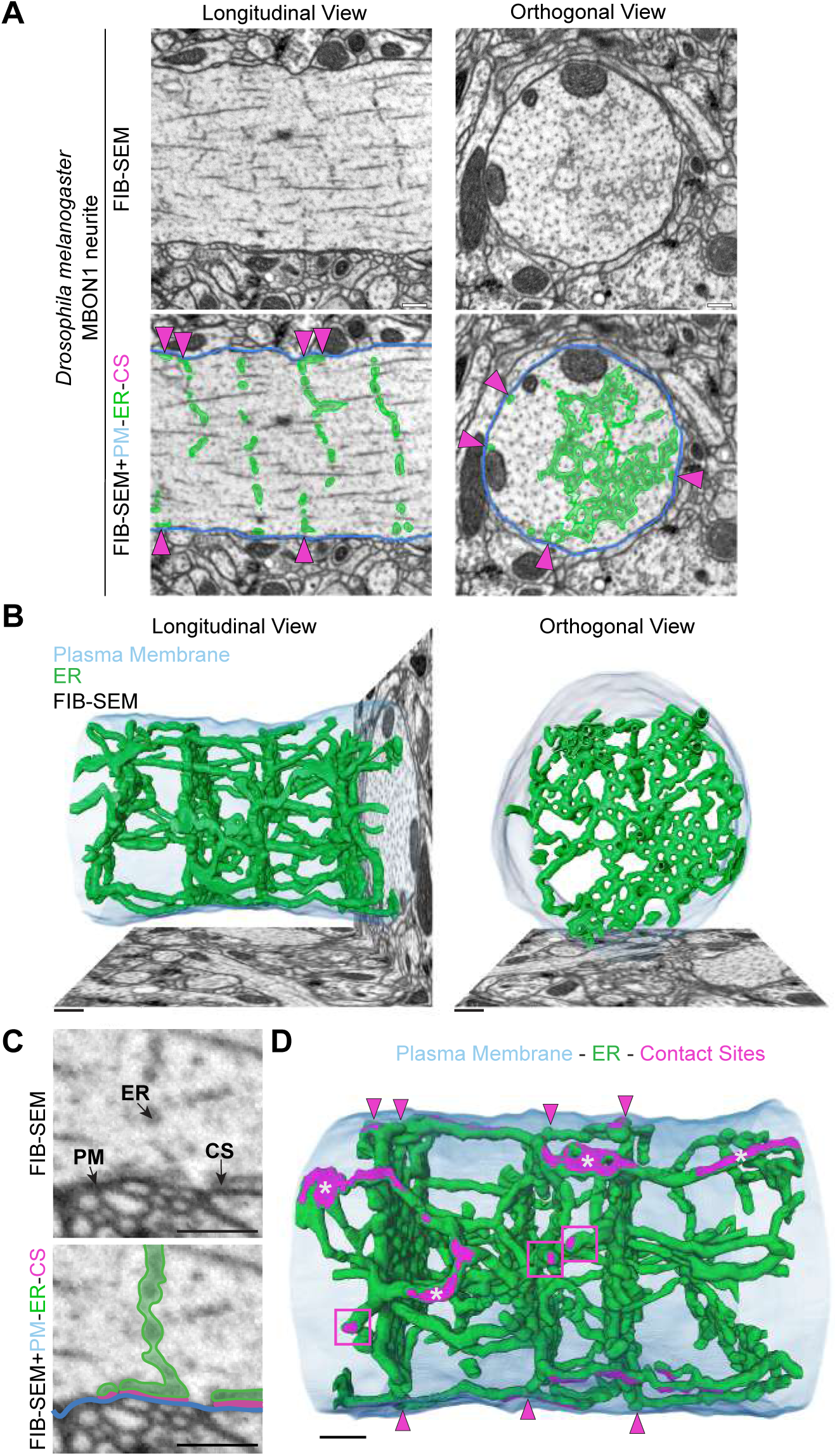
The ladder-like array organization of the ER is a conserved feature in invertebrate neurons. A) FIB-SEM orthoslices showing the morphology of the ER in longitudinal and orthogonal sections of the fly MBON1 neurite (top). The segmentation of endoplasmic reticulum (ER, green) and plasma membrane (PM, blue) are shown (bottom). Junctional sites between ER and PM visible in the orthoslices are indicated with magenta arrowheads. Scale bars: 1 μm. B) Longitudinal and orthogonal views of the 3D renderings of ER (green) and plasma membrane (blue) in a MBON1 neuron segmented from FIB-SEM datasets. Scale bars: 1 μm. C) *Drosophila* ER-PM junction as visualized in a single FIB-SEM orthoslice (top panel) with overlay of endoplasmic reticulum (ER, green), plasma membrane (PM, blue) and contact sites (CS, magenta) (bottom panel). Scale bar: 250 nm. D) 3D renderings of ER (green), plasma membrane (blue) and ER-PM contact sites (magenta) in fly MBON1 neuron segmented from FIB-SEM datasets. Magenta arrowheads and square boxes highlight contact sites anchoring ER arrays to the plasma membrane. Asterisks indicate non-rung associated contact sites. Scale bar: 1 μm.

### Stability and dynamism of ER and ER-PM junctions in dendrites

We next explored the stability and dynamism of laddered ER and ER-PM junctions in dendrites in rat hippocampal neurons. Using 2D lattice-SIM imaging, we acquired time-lapse sequences at 20 ms intervals. ER-PM junctions were found to be stable over minutes, while laddered ER elements displayed dynamic behavior, constantly protruding, retracting, narrowing, or expanding their caliber over millisecond to second timescales (Figure 3A, Movie 3). To test whether ER-PM junctions serve to anchor laddered ER elements, we exposed neurons to hypotonic stress, which causes the ER to vesiculate into giant vesicles without disrupting ER-PM contact sites^47^. After such treatment, dendritic ER ladders transformed into giant ER vesicles. The vesicles were distributed at regular intervals throughout the dendrite, as visualized by the PM marker mTagBFP2-CAAX2, and they maintained contact with the PM at JPH3-enriched ER-PM junctions, as shown using mCherry-JPH3 (Figure 3B). Notably, mCherry-JPH3 puncta remained regularly distributed along dendrites at the same interval of ∼1 μm (0.9 ± 0.1 μm, Figure 3C), as seen prior to hypotonic stress. This suggested that ER-PM junctions anchor and stabilize ER elements to the PM even when these ER elements are dramatically perturbed during hypotonic stress.

**Figure 3:**
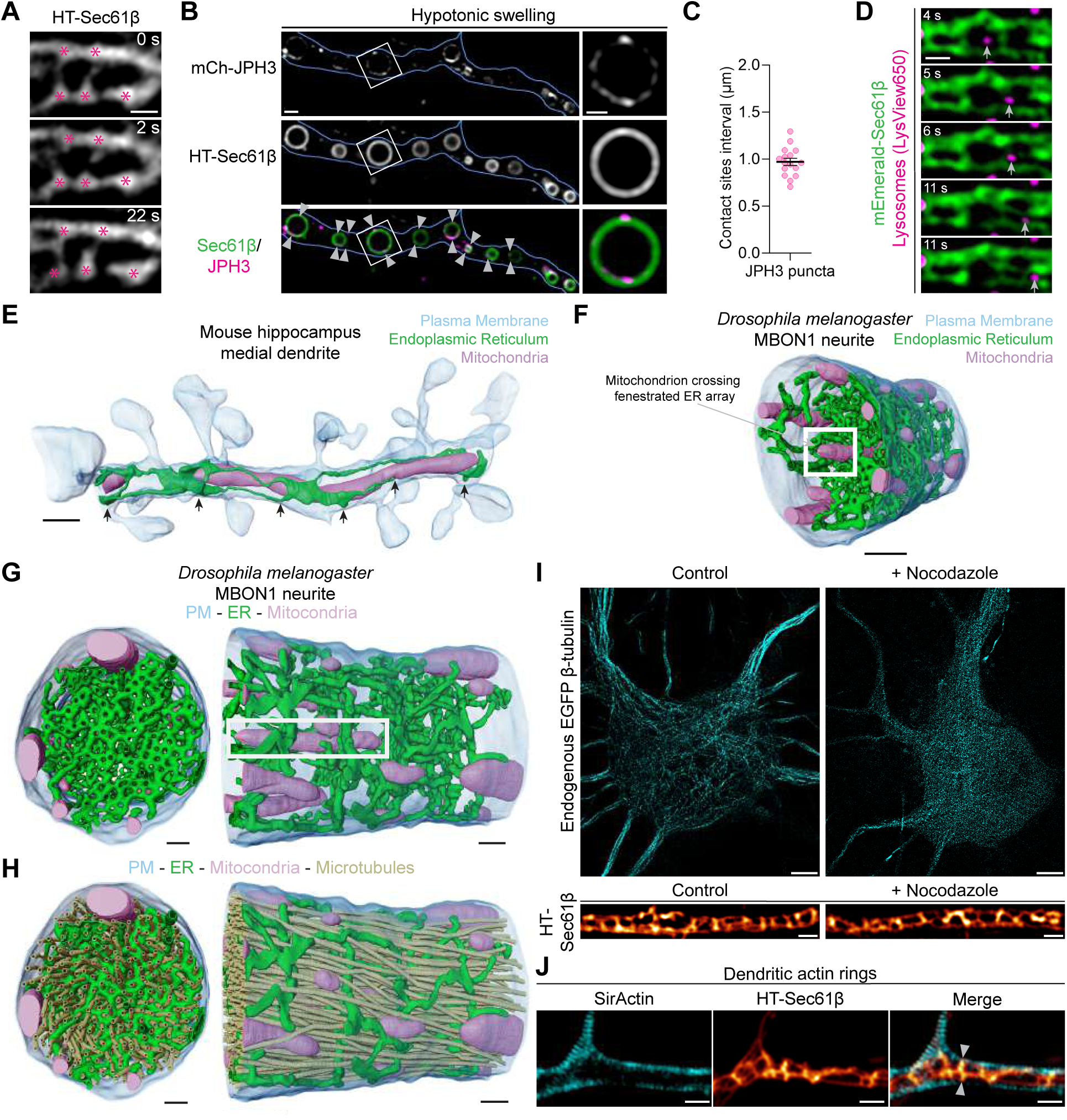
Regularly distributed ER-PM junctions persist under major cytoskeletal perturbations. A) Time-lapse acquired using 2D lattice-SIM in burst mode of HaloTag-Sec61β (labeled with JF585 HaloTag-ligand) expressing neurons revealing ER tubule dynamicity with the persistent presence of ER-PM junctional sites (marked with magenta asterisks, see Movie 3). Scale bar: 0.5 μm. B) Neuronal dendrite expressing mCherry-JPH3, mEmerald-Sec61β and mTagBFP2-CAAX2 (blue outline) following hypotonic swelling (∼63 mOsM solution). Scale bar: 1 μm, inset 0.5 μm. C) Average mCherry-JPH3 contact sites interval after treatment with strong hypotonic solution (0.97 ± 0.15 μm, n = 16, N =2, mean ± SD). D) Time-lapse series acquired using Lattice-SIM of mEmerald-Sec61β expressing neurons. Lysosomes were stained using Lysview650. Lysosomes move through ladder-like ER arrays (see arrow and Movie 4). Scale bar: 0.5 μm. E) 3D renderings of ER (green), plasma membrane (blue) and mitochondria (pink) in medial dendrites of mouse hippocampus. Black arrows indicate ER rungs. Scale bar: 1 μm. F-G) 3D renderings of ER (green), plasma membrane (blue) and mitochondria (pink) in *Drosophila* MBON1 neuron segmented from FIB-SEM datasets. The white box frames a mitochondrion extending through multiple ER rungs. Scale bars: 1 μm. H) Orthogonal and longitudinal views of the 3D renderings of ER (green), plasma membrane (blue) mitochondria (pink) and microtubules (tan) in *Drosophila* MBON1 neuron segmented from FIB-SEM datasets. Scale bars: 1 μm. I) ) Primary hippocampal neurons were transiently transfected with pORANGE Tubb3-GFP KI, to endogenously tag tubulin, allowing visualization of the microtubule cytoskeleton. Images show microtubule organization in untreated cells (Control) and nocodazole treated cells (5 μM nocodazole for 2-3 h, microtubule destabilization is evident in cell bodies and proximal dendrites). Even after prolonged nocodazole-induced destabilization of the microtubule cytoskeleton, the laddered ER organization remained unchanged. The ER was imaged in the same neurons also expressing HaloTag-Sec61β (labeled with JF585 HaloTag-ligand). Scale bar: 5 μm in cell body and 0.5 μm in dendrites. J) Lattice-SIM image of dendritic actin rings (SirActin labeling) and ladder-like ER (HaloTag-Sec61β labeled with JF549 HaloTag-ligand). Arrowheads show ER tubules protruding between dendritic actin rings to establish contacts with the PM. Scale bar: 1 μm.

Laddered ER in dendrites were not a barrier to motile organelles as demonstrated by lysosomes moving along dendrites. The lysosomes readily trafficked through the rungs of laddered ER and while doing so, transiently altered ER ladder organization but not ER-PM junction arrangement (Figure 3D, Movie 4). Mitochondria, too, appeared to be capable of penetrating the dense array of tubules in the rungs of laddered ER, as shown in the FIB-SEM dataset of a single mitochondria spanning the center of several consecutive ER rungs both in mouse (Figure 3E) and fly neurites (Figure 3F and Figure 3G). In addition to lysosomes and mitochondria freely passing through ER rungs, FIB-SEM datasets showed microtubules extending through fenestrae and holes of ER rungs (Figure 3H, Figure S3C), indicating that ER rungs are not barriers to these structures.

The maintenance of laddered ER organization in dendrites was independent of microtubules since microtubule depolymerization using 5 µM nocodazole for 2-3 h neither disrupted the laddered ER morphology nor perturbed periodic contacts of the ER with the PM (Figure 3I). Laddered ER organization also appeared to be independent of actin, at least for periodic actin rings (Figure 3J,), which circumscribe the PM of dendrites^48^. The periodicity of actin rings (∼190 nm)^49^ was much smaller than the periodicity of ER ladders (∼1 µm), and ER tubules protruded between actin rings to reach ER-PM contact sites (Figure 3J, arrowheads). ER tubule association with ER-PM junctions did not require actin as ER-PM junctions and ER remained associated under hypotonic stress (see Figure 3B), which causes actin disassembly.

### Junctophilin-3 coordinates the recruitment of voltage-gated calcium channels to ER-PM junctions

Two PM-localized voltage-gated Ca^2+^ channels, Ca_V_2.1 and Ca_V_2.2, concentrate at ER-PM junctions in non-neuronal cells and their inactivation is slowed when JPH3 is overexpressed^50^. This observation led us to ask whether the regularly spaced, JPH3-positive ER-PM junctions we observed in dendrites were hubs of Ca^2+^ release and uptake machinery. To address this question, we used modified lentiviral vectors (see Methods) to express mEmerald-Ca_V_2.1 or mEmerald-Ca_V_2.2 in cultured primary rat hippocampal neurons. Live-cell imaging experiments revealed mEmerald-Ca_V_2.1 and mEmerald-Ca_V_2.2 to be present in nerve terminals (Figure S4A), neuronal cell bodies (Figure S4B and S4C) and dendrites (Figure 4A and 4B, Figure S4A). In dendrites, Ca_V_2.1 and Ca_V_2.2 resided in small puncta distributed at roughly regular intervals across the PM (Figure 4A and 4B). These puncta resided at sites where ER rungs intersected the PM based on antibody labeling for Ca_V_2.1 and an ER marker (Figure S4D).

**Figure 4:**
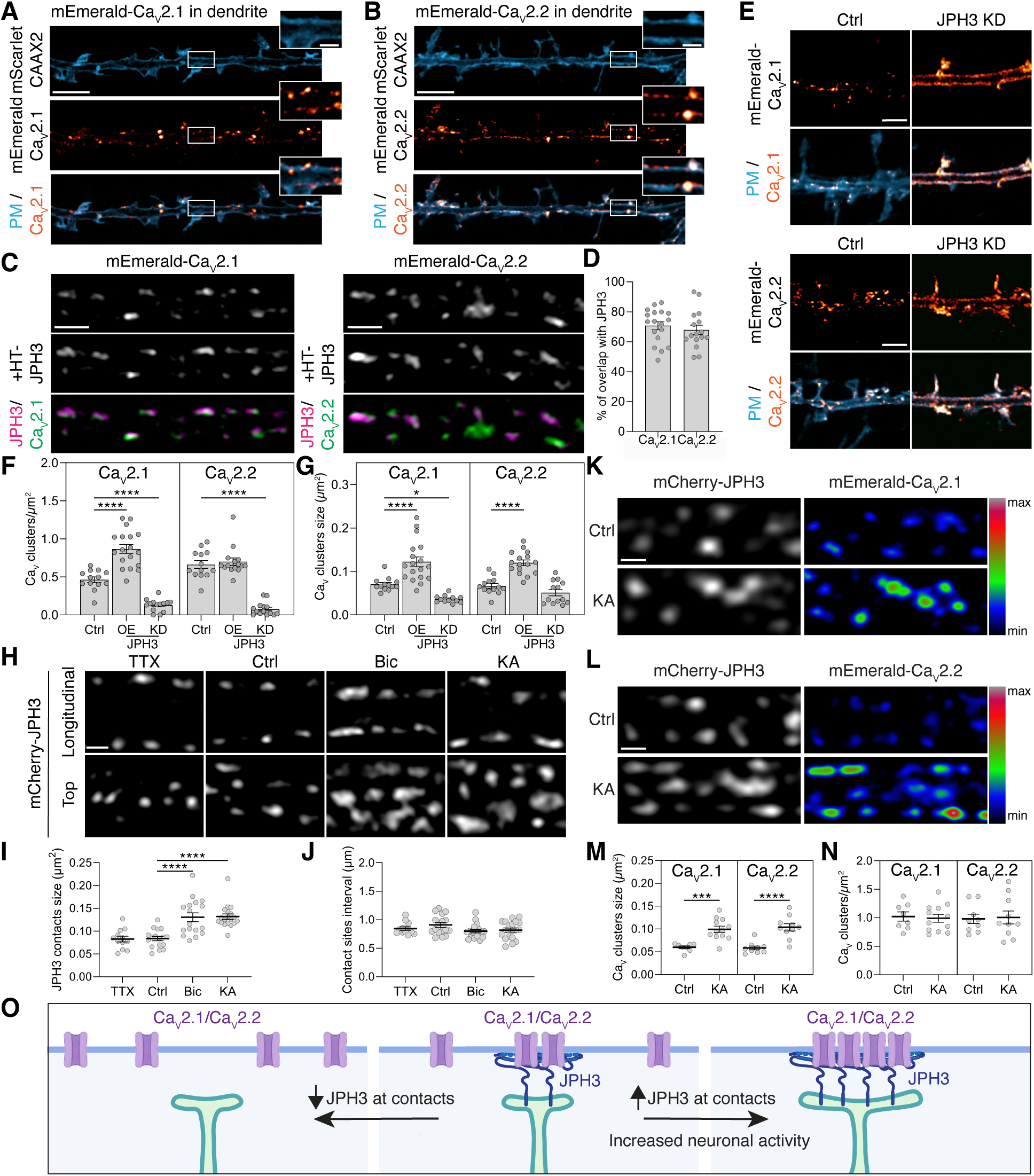
Junctophilin-3 coordinates the recruitment of voltage-gated calcium channels to ER-PM contact sites. A-B) Airyscan images showing mEmerald-CaV2.1 (A) and mEmerald-CaV2.2 (B) localize at the plasma membrane (visualized with mScarlet-CAAX2) of dendrites in primary rat hippocampal neurons. mEmerald-CaV2.1 and mEmerald-CaV2.2 accumulated in regularly interspaced puncta (inset). Scale bar: 5 μm, inset: 1 μm. C) mEmerald-CaV2.1 and mEmerald-CaV2.2 colocalized with HaloTag-JPH3 contacts in neuronal dendrites. Scale bar: 1 μm. D) mEmerald-CaV2.1 and mEmerald-CaV2.2 puncta were observed in approximately 70.7 ± 2.7% (n =18, N= 5) and 68.0 ± 3.2% (n =16, N= 4) of JPH3-positive contacts, respectively, along the dendritic tree. E) Knockdown of JPH3 resulted in the loss of mEmerald-CaV2.1 and mEmerald-CaV2.2 clusters in dendrites. Neurons expressing shRNA for JPH3 were identified by cytosolic expression of mTagBFP2. Scale bars: 2 μm. F) mEmerald-CaV2.1 hotspots at contact sites per dendritic surface unit was approximately 0.46 ± 0.04 clusters/μm2 (n=13, N=5). Co-overexpression with HaloTag-JPH3 significantly increased the density of mEmerald-CaV2.1 clusters (0.87 ± 0.06%, n = 18, N = 5) while JPH3 knock down significantly reduced mEmerald-CaV2.1 clusters density (0.13 ± 0.02 clusters/μm2, n = 14, N = 4). The density of mEmerald-CaV2.2 was 0.66 ± 0.05 clusters/μm2 (n=13, N=4). While the co-overexpression of HaloTag-JPH3 did not significantly increase the density of mEmerald-CaV2.2 clusters (0.70 ± 0.05 clusters/μm2, n = 15, N = 4); JPH3 knock down significantly decreased mEmerald-CaV2.2 clusters density (0.08 ± 0.02 clusters/μm2, n = 14, N = 3). G) HaloTag-JPH3 significantly increased the size of mEmerald-CaV2.1 and mEmerald-CaV2.2 clusters at contact sites (mEmerald-CaV2.1 = 0.070 ± 0.004 μm2, n = 13, N =5; mEmerald-CaV2.1 + JPH3 OE = 0.122 ± 0.011 μm2, n = 18, N = 5; mEmerald-CaV2.1 + JPH3 KD = 0.036 ± 0.002, n=12, N =4; mEmerald-CaV2.2 = 0.067 ± 0.005, n = 13, N = 4; mEmerald-CaV2.2 + JPH3 OE = 0.120 ± 0.006 μm2, n = 16, N =4; mEmerald-CaV2.2 + JPH3 KD = 0.051 ± 0.007 μm2, n=13, N =3). H) Morphology and distribution of HaloTag-JPH3 junctions in neuronal dendrites after 12-hour treatment with TTX (0.5 μM), Bicuculline (Bic, 20 μM), and Kainic Acid (KA, 250 nM). Scale bar: 1 μm. I) HaloTag-JPH3 contact site sizes in Ctrl conditions (0.084 ± 0.005 μm2, n = 17) and after treatments with TTX (0.082 ± 0.006 μm2, n = 12), Bic (0.131 ± 0.009 μm2, n = 17), and KA (0.132 ± 0.006 μm2, n = 19); N =3. J) Comparison of HaloTag-JPH3 contact site intervals in control (Ctrl) conditions (0.91 ± 0.04 μm, n = 17) and after treatments with TTX (0.85 ± 0.03 μm, n = 12), Bic (0.80 ± 0.03 μm, n = 17), and KA (0.82 ± 0.04 μm, n = 19); N = 3. K-L) mEmerald-CaV2.1 (K) and mEmerald-CaV2.2 (L) plasma membrane puncta colocalizing with mCherry-JPH3 increased size upon KA treatment. Scale bars: 1 μm. M) mEmerald-CaV2.1 clusters size at contacts in Ctrl (0.060 ± 0.003 μm2, n = 8) and after KA incubation (0.099 ± 0.007 μm2, n = 12), mEmerald-CaV2.2 clusters size at contacts in Ctrl (0.058 ± 0.004 μm2, n = 9) and after KA incubation (0.103 ± 0.008 μm2, n = 10); N =3. N) Comparison of mEmerald-CaV2.1 clusters density in control (Ctrl) conditions (1.02 ± 0.08 clusters/ μm2, n = 8) and after KA treatment (0.99 ± 0.07 clusters/μm2, n = 12) and mEmerald-CaV2.2 clusters density in control (Ctrl) conditions (0.98 ± 0.08 clusters/μm2, n = 9) and upon KA treatment (1.01 ± 0.11 clusters/μm2, n = 10); N = 3. O) Schematic illustrating how changes in ER-PM contact site size and modulation of JPH3 accumulation affect voltage-dependent Ca2+ channel subdomains on the plasma membrane. n = neurons, N = independent experiments. All data are mean ± SEM and were analyzed with one-way ANOVA and Tuckey’s multiple comparison tests; * p<0.05, *** p<0.001, **** p<0.0001.

Co-expression of HaloTag-JPH3 with mEmerald-Ca_V_2.1 and mEmerald-Ca_V_2.2 revealed a robust colocalization of Ca_V_2.1 or Ca_V_2.2 in the same puncta as JPH3 in both dendrites and cell bodies (Figure 4C and Figure S4E). Along the dendritic tree, JPH3-positive puncta were coincident with Ca_V_2.1 (70%) and Ca_V_2.2 (68%) (Figure 4D). Upon JPH3 overexpression conditions, the number of Ca_V_2.1 and Ca_V_2.2 clusters per unit dendritic-area and their size significantly increased (Figure 4F, 4G and Figure S4E), while knockdown of JPH3 (Figure 4E and Figure S4F-H) led to a significant reduction in Ca_V_2.1 and Ca_V_2.2 clusters number and size at ER-PM junctional sites in dendrites (Figure 4E-G) and cell bodies (Figure S4H). These findings suggest that JPH3’s presence at neuronal ER-PM junctions in neurons determines the extent to which Ca_V_2.1 and Ca_V_2.2 accumulate at these sites (Figure 4O).

To address whether Ca_V_2.1/ Ca_V_2.2 containing, JPH3-positive ER-PM junctions are responsive to changes in neuronal activity, we compared HaloTag-JPH3 contact site size and interval in neuronal dendrites following treatment with inhibitors and activators of neuron excitability. A 12 h treatment with the voltage-gated Na^+^ channel blocker tetrodotoxin (TTX) was used to decrease excitability, while the GABA_A_ antagonist, bicuculline (Bic), or the glutamate receptor agonist, kainic acid (KA), enhanced excitability (Figure 4H-J). Increasing excitability increased the size of JPH3-labeled contact sites (Figure 4H-I) and the abundance of Ca_V_2.1 and Ca_V_2.2 at contact sites (Figure 4K-M) without affecting contact site intervals or density (Figure 4J and 4N). Therefore, increases in neuronal activity enhance the quantity of both Ca^2+^ channels and JPH3 proteins at ER-PM contact sites (Figure 4O).

### ER-PM contacts represent sites for pCaMKII-dependent intracellular signaling

The calcium-calmodulin-dependent protein kinase II (CaMKII) is activated by the association with the Ca^2+^-calmodulin complex upon elevation in intracellular [Ca^2+^]. This binding leads to CaMKII autophosphorylation (pCaMKII), which then phosphorylates various proteins, altering their structure and functions to trigger molecular pathways required for synaptic plasticity^51,52^. Given the localization of Ca_V_2.1 and Ca_V_2.2 channels to JPH3-positive ER-PM junctions, we tested whether ER-PM contacts serve as sites for the initiation of pCaMKII-dependent intracellular signaling. We employed a depolarizing paradigm that maximizes the contribution of voltage-gated Ca^2+^ channels to depolarization-induced Ca^2+^ entry^20^ (Figure 5A). Upon depolarizing rat primary hippocampal neurons with 90 mM K^+^ for 10 s, focal accumulations of pCaMKII were seen both in spine heads and all along dendrites^53^ (Figure 5B, central panels, Figure 5C). This pCaMKII accumulation occurred without affecting the frequency of JPH3-positive contact sites (Figure 5D). Localizing the α isoform of CaMKII (CaMKIIα) before and after depolarization by immunofluorescence revealed that CaMKIIα accumulated at ER-PM junctions after PM depolarization (Figure S5A). Notably, pCaMKII hotspots colocalized with HaloTag-JPH3 positive contact sites along the dendritic shaft (Figure 5B central panels, Figure 5E). Incubation with Ca_V_ blockers (the L-type Ca_V_ blocker nimodipine, the P/Q-type Ca_V_ blocker **ω**-Agatoxin IVA, and N-type Ca_V_ blocker **ω**-Conotoxin GVIA ) effectively prevented pCaMKII accumulation at ER-PM contact sites (Figure 5B bottom panels, Figure 5C and 5E), suggesting that depolarization-induced opening of Ca_V_ channels increased [Ca^2+^] sufficiently at the ER-PM contacts to stimulate pCaMKII localization, and their inferred activity. Similar results were seen in neuronal cell bodies, suggesting that ER-PM contact sites represent a location of pCaMKII accumulation also in cell bodies (Figure S5B and S5C).

**Figure 5:**
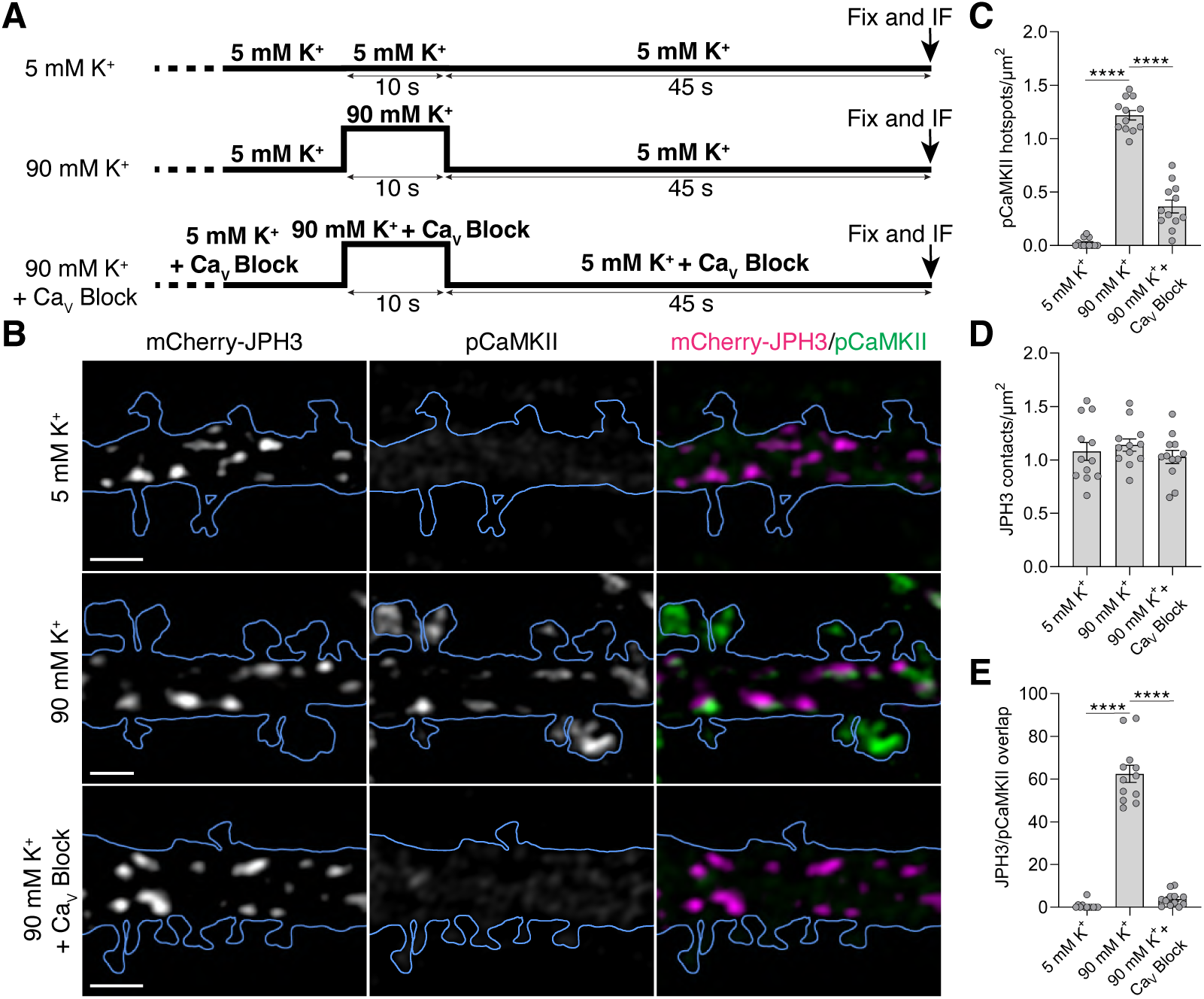
pCaMKII accumulates at ER-PM junctions upon neuronal stimulation. A) Protocols used to detect pCaMKII and CaMKIIα at ER-PM junctions using a paradigm that maximizes the contribution of voltage-gated Ca^2+^ channels to depolarization-induced Ca^2+^ entry. B) Dendrites expressing mTagBFP2-CAAX2 to visualize dendritic plasma membrane (blue outline) and mCherry-JPH3 to visualize ER-PM junctions were promptly fixed and stained for pCaMKII under different conditions: 5 mM K^+^, stimulation with 90 mM K^+^, and 90 mM K^+^ with Ca_V_ blockers: the L-type Ca_V_ blocker nimodipine (10 μM), the P/Q-type Ca_V_ blocker **ω**-Agatoxin IVA (500 nM), and **ω**-Conotoxin GVIA (2 μM). The background signal of pCaMKII staining was subtracted to highlight hotspots (see Methods section). Scale bars: 1 μm. C) pCaMKII hotspots per μm^2^ of dendritic area in 5 mM K^+^ = 0.02 ± 0.01, 90 mM K^+^ = 1.2 ± 0.04%, 90 mM K^+^ + CaV Block = 0.36 ± 0.06 hotspots/μm^2^, n=12, N=3. D) JPH3 contacts density in dendrites in each stimulation paradigm: 5 mM K^+^ = 1.08 ± 0.08 μm^2^, 90 mM K^+^ = 1.14 ± 0.06, 90 mM K^+^ + Ca_V_ Block = 1.03 ± 0.06 contacts/μm^2^, n=12, N=3. E) Fraction of HaloTag-JPH3 contacts colocalizing with pCaMKII hotspots in 5 mM K^+^ = 0.67 ± 0.49%, 90 mM K^+^ = 62.44 ± 4.02%, 90 mM K^+^ + Ca_V_ Block = 3.88 ± 0.98%, n=12, N=3. All data are mean ± SEM and were analyzed with one-way ANOVA and Tuckey’s multiple comparison tests; **** p<0.0001.

### Dendritic ER-PM contacts are populated by a machinery for the fine-tuning of ER Ca^2+^ uptake and release

We next explored the distribution of ER-associated Ca^2+^ release and uptake machinery in dendrites, asking whether any component of this machinery was localized at JPH3-positive ER-PM junctions. The machinery we examined included stromal interaction molecules 1 and 2^54–58^ (STIM1 and STIM2, ER-localized calcium sensors that activate Orai Ca^2+^ ion channels on the PM), ER Ca^2+^ATPase^59^ (SERCA, the pump that transfers Ca^2+^ from cytosol into ER lumen), and the ryanodine receptors^50,60^ (ER Ca^2+^ release channels).

Examination of primary neurons expressing HaloTag-STIM1 or HaloTag-STIM2 together with an ER marker under basal conditions revealed that STIM2 was enriched at ER-PM junctions (Figure 6A and Figure S6A), whereas STIM1 was distributed diffusely on ER ladders (Figure 6B and Figure S6B) and populated ER-PM junctions upon ER Ca^2+^ depletion (Figure S6C).

**Figure 6:**
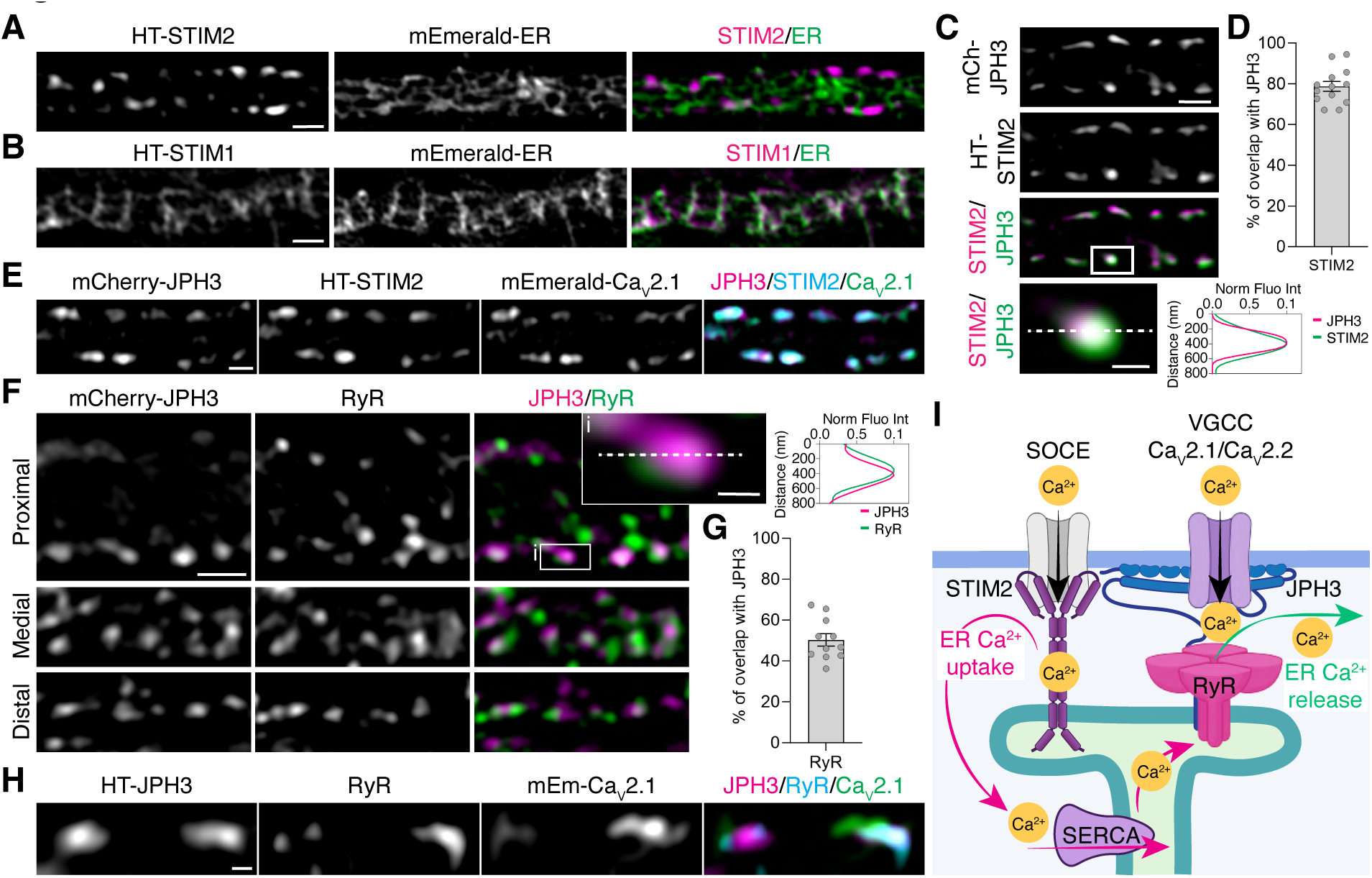
R**e**gular **ER-PM contact sites host ER Ca^2+^ uptake and release machinery.** A) Dendrite expressing HaloTag-STIM2 with the ER marker mEmerald-ER. Halo-STIM2 accumulated at regularly distributed contacts in dendrites. Scale bar: 2 μm. B) Primary neuronal dendrite co-expressing HaloTag-STIM1 with the ER marker mEmerald-ER showed Halo-STIM1 distribution within ER rungs. Scale bar: 2 μm. C) Halo-STIM2 colocalized with regularly spaced mCherry-JPH3 contacts in neuronal dendrites. Inset and fluorescence intensity plot profiles emphasize the overlap between JPH3 hotspots and STIM2 clusters. Scale bar: 1 μm main image, 0.25 μm inset. D) HaloTag-STIM2 populated ∼78.7 ± 2.4% of JPH3-positive contacts in neuronal dendrites (n =13, N= 3). E) Co-overexpression of mCherry-JPH3, HaloTag-STIM2 and mEmerald-Ca_V_2.1 showed the colocalization of these molecules at regular ER-PM dendritic contacts. Scale bar: 0.5 μm. F) Immunofluorescence experiments revealed the colocalization of RyRs puncta with JPH3-positive ER-PM junctions throughout proximal, medial, and distal dendrites. Inset and fluorescence intensity plot profiles emphasize the overlap between JPH3 and RyRs clusters. Scale bar: 1 μm main image, 0.25 μm inset. G) RyRs puncta populated ∼50.3 ± 3.1% of JPH3 contacts in dendrites (n =11, N= 2, mean ± SEM). H) Immunofluorescence experiments revealed the colocalization of RyRs, HaloTag-JPH3, and mEmerald-Ca_V_2.1 at periodic ER-PM contacts in dendrites. Scale bar: 0.2 μm. I) Schematic illustrating the molecular components of ER Ca^2+^ uptake and release machinery populating regularly distributed ER-PM contacts in dendrites. Data are reported as mean ± SEM. Constructs fused to HaloTag were labeled with JF635 or JF647.

The STIM2 puncta at ER-PM junctions seen under basal conditions were distributed at the same regular intervals found for JPH3 (Figure S6D and S6E), with 82% of JPH3-positive puncta populated with STIM2 (Figure 6C and 6D). ER-PM junctions populated by JPH3 and STIM2 colocalize with the CaV2.1 hotspots at the plasma membrane (Figure 6E and zoom-in in Figure S6F). In contrast to STIM2, SERCA, examined by immunofluorescent labeling, was uniformly distributed across the ER ladder-like array in dendrites, with no accumulation at ER-PM junctions (Figure S6G).

To examine ryanodine receptors (RyRs) distribution, we conducted immunofluorescence experiments using a pan anti-RyRs antibody (that recognized all the different isoforms of the channel), in rat primary neurons expressing a fluorescent marker to visualize the ER (HT-Sec61β) and mCherry-JPH3. RyRs puncta colocalized with JPH3-positive ER-PM junctions in ER in proximal, medial, and distal zones of dendrites (Figure 6F), with RyRs present in approximately 50% of JPH3-positive contacts (Figure 6G), often at both extremities of the ER rungs (Figure S6H). Together, these observations indicated that the ER at JPH3-positive ER-PM junctions are basally populated by specific types of Ca^2+^ release and uptake machinery, including STIM2 and RyRs, with SERCA and STIM1 more widely distributed. The presence of ER-associated Ca^2+^ handling machinery (i.e., STIM2 and RyRs) together with the PM-associated voltage gated Ca_V_2.1 and Ca_V_2.2 channels at JPH3-positive ER-PM junctions (Figure S6F, Figure 6H and zoom-in in Figure S6I and Figure 6I) raised the question of whether these periodically aligned junctions serve as hubs for serial Ca^2+^ flux and signaling along dendrites.

### Ryanodine-dependent ER Ca^2+^ release occurs at regularly defined sites and can be detected more than 20 μm away from individual spines upon stimulation

To address the above question regarding the role of periodically aligned ER-PM junctions, we examined whether focal synaptic inputs, which lead to exponentially decaying cytosolic Ca^2+^ signals^61^, might, in turn, drive activation of RyRs at ER-PM junctions distal from the given synaptic input, which, in turn, might amplify the local Ca^2+^ concentration and recruit Ca^2+^-dependent proteins such as pCaMKII to these sites. To investigate this possibility, we expressed the ER calcium indicator ER-GCaMP6-210^62^, along with the post-synaptic marker PSD95-mCherry or the cytosolic Ca^2+^ sensor RCaMP1.07 in primary hippocampal neurons (Figure 7A). We then used two-photon glutamate uncaging to provide focal spine activation capable of eliciting measurable signal changes in ER and cytosolic Ca^2+^ and measured both local and distal Ca^2+^ levels.

**Figure 7:**
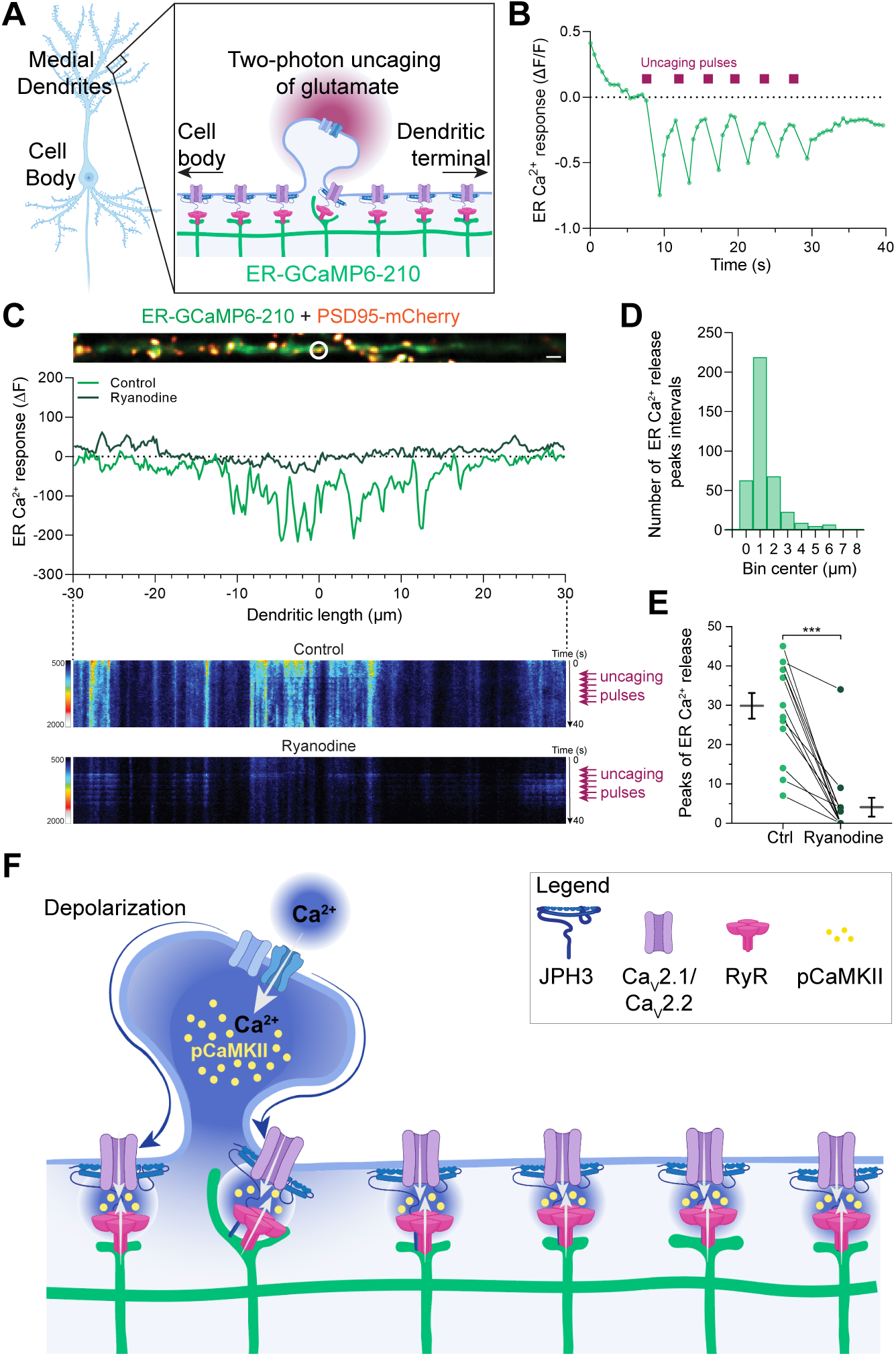
Regularly distributed ER-PM contacts allow RyR-dependent long-distance propagation of Ca^2+^ signals in dendrites. A) Illustration communicating the experiments monitoring ER Ca^2+^ dynamics in dendrites localized ∼70 μm away from the cell body of primary hippocampal neurons expressing ER-GCaMP6-210 upon two-photon glutamate uncaging. B) Representative temporal profile of dendritic ER Ca^2+^ release following 6 pulses of glutamate uncaging. The burgundy squares indicate the glutamate uncaging events. C) Dendrite expressing ER-GCaMP6-210 and PSD95-mCherry. The average plot and the Control and Ryanodine kymographs illustrate the ER Ca^2+^ profile measured in the same dendrite before, during, and after the application of the RyRs inhibitor ryanodine. Scale bar: 2 μm. D) Frequency distribution of ER Ca^2+^ release intervals (n = 404 intervals, n = 14 dendrites, N=3 biological replicates. E) Impact of Ryanodine treatment on the reduction of ER Ca^2+^ release events from dendritic ER upon local spine stimulation. (Control: peaks n = 29.86 ± 3.2, Ryanodine: peaks n = 4.07 ± 2.4; across 14 dendrites, N=3 biological replicates). Data are reported as mean ± SEM. *** p<0.001. F) Illustration showing how stimuli originating from a single spine could impact local and distal membrane proximal biochemistry at sites tens of μm away along a dendrite through serial activation of Ca^2+^ release and uptake machinery at periodic ER-PM junctions. Arrows denote Ca^2+^ flux.

Repeated uncaging at the same spine led to robust abrupt changes in ER Ca^2+^ in 48% of dendrites examined, peaking at ∼ 680 msec after the uncaging event and recovering over the next few seconds (Figure 7B and Figure S7A). The average line scan signal across many dendrites showed that focal uncaging leads to a net decrease in ER Ca^2+^ in the vicinity of the activated spine (Figure S7B). The magnitude of the drop decayed laterally away from the activation site, similar to the spatial decay of the signal in the cytoplasm (Figure S7B). RyRs at dendritic ER-PM junctions are involved in this ER Ca^2+^ efflux, as the drop in ER Ca^2+^ was eliminated upon blocking RyRs with the drug ryanodine, resulting in a small net elevation in ER Ca^2+^ upon spine activation (Figure S7C).

Line scans of ER Ca^2+^ signal from individual dendrites showed a highly structured spatial signal (Figure 7C, Figure S7D) suggestive of ER Ca^2+^ release at discrete intervals distal to the activated spine. Kymographs of the ER Ca^2+^ images before, during and after each uncaging pulse (Figure 7C, Control kymograph) revealed that the ER Ca^2+^ release driven by spine stimulation was spatially structured (Figure 7C, Control kymograph), with average individual peaks in the data appearing at ∼ 1 μm intervals (Figure 7D and Figure S7D), similar to the spacing of ER-PM junctions (Figure 1), and over 20 μm away in either directions (Figure 7C and Figure S7D and S7E).

Notably, all peaks disappeared upon stimulation in the presence of the ryanodine receptor inhibitor ryanodine (Figure 7C, Ryanodine kymograph, and Figure 7E). Trial-to-trial examination of the line scans showed that although the response at individual locations along the dendrite could fluctuate across trials, many persisted throughout (Figure S7E). The results thus support the idea that ER-PM junctions serve as the structural element supporting Ca^2+^ release from the ER. In this view, stimuli originating at one spine location could impact local membrane biochemistry at sites tens of μm away in the dendrite through serial activation of Ca^2+^ release and uptake machinery at periodic ER-PM junctions (Figure 7E).

## Discussion

A major question in neuronal cell biology has been how postsynaptic neurotransmission is integrated over long distances in dendrites^63^. Ca^2+^ is a pivotal second messenger in this process, with its cytoplasmic transients tightly controlled both spatially and temporally by buffers, pumps, exchangers, and other Ca^2+^ binding proteins^3^. Prior imaging studies have suggested that synaptic receptor-dependent Ca^2+^ transients are mostly confined within spine heads, with only minimal spreading along dendritic shafts^4,64–66^. This suggests alternative mechanism(s) to simple spine-based Ca^2+^ release to support long-range Ca^2+^-dependent intracellular responses in dendrites.

Mechanisms previously proposed to accomplish this include 1) local increases in Ca^2+^ within spines activating transport and subsequent translocation of Ca^2+^-regulated proteins from their synaptic sites of activation, 2) influx of Ca^2+^ through somatic and perisomatic voltage-gated Ca^2+^ channels upon depolarization, 3) propagation of inositol trisphosphate receptor (IP3R)-dependent waves upon release of Ca^2+^ from internal stores; and 4) passive propagation of electrical potentials along ER membranes (reviewed in ^67^).

Here, we report an additional mechanism that supports long-range Ca^2+^ responses in dendrites involving a specialized organization of ER and PM. We found that neuronal dendrites organize their ER and PM to form regularly arrayed junctions enriched in Ca^2+^ uptake and release machinery. The ER-PM junctions are arranged at roughly 1-µm intervals throughout the entire dendritic tree, with short transversal ER elements intersecting the PM at perpendicular angles and exhibiting a ladder-like morphology in proximal and medial dendrites. Populated by RyRs and voltage-gated Ca^2+^ channels and tethered in part by JPH3, the ER-PM junctions in dendrites bear similarities to such junctions in skeletal, cardiac, and smooth muscle ^39,40,68,69^ as well as neuronal subsurface cisternae^7^. We showed that ER-PM contacts increase the signaling capacity of a dendrite beyond the simple excitation of a propagating action potential. By utilizing ER Ca^2+^ release at micron intervals, calcium signals by themselves may be propagated as they do in regenerative Ca^2+^ waves in the soma of many cells activated by Gq-linked G protein-coupled receptors (GPCRs) with a mechanism that distributes PM Ca^2+^ entry through much larger volumes^70^. An important area to be studied is whether dendritic Gq-linked GPCRs might modify the ER Ca^2+^ store or induce release on their own, possibly disrupting or enhancing signals from the synapse. Furthermore, a dendritic signal may be enriched by the range of consequential responses, such as during moderate changes in synaptic calcium that might be propagated in the absence of an action potential or during afterhyperpolarization events that follow bursts of action potentials often due to the activation of Ca^2+-^dependent potassium currents^28,71^.

Consistent with a long-distance intracellular Ca^2+^ signal integration role of ER-PM interplay in dendrites^7,17,19–21,61^, we found that the periodic ER-PM junctions coordinated Ca^2+^-induced Ca^2+^ release by RyRs over 20 µm away from locally stimulated spines with Ca^2+^ levels in ER changing across a spatial scale ∼100 times greater than that of the spine neck. In this mechanism, ER-PM junctions arranged at ∼1 µm intervals through the neuron-specific ER-PM tethering protein JPH3 coordinates the recruitment of voltage-gated Ca^2+^ channels (specifically Ca_V_2.1 and Ca_V_2.2). This arrangement creates a repeating microenvironment where Ca^2+^ entry from Ca_V_2.1 and Ca_V_2.2 can reach the necessary micromolar range for activation of Ca^2+^-releasing ryanodine receptors on ER. These Ca^2+^ transients are also sufficient for calmodulin-dependent phosphorylation of CaMKII at ER-PM contacts. Supporting this mechanism, we found that alterations in JPH3 accumulation at ER-PM junctions resulted in changes in voltage-gated Ca^2+^ channel cluster size and its knockdown in the dispersal of Ca_V_2.1 and Ca_V_2.2 subdomains at the PM. Constitutive localization of STIM2 at the ER-PM junctions may also facilitate rapid Ca^2+^ entry from the plasma membrane, with ER refilling occurring from more broadly distributed SERCA pumps (see the illustrations in Figure 6I and Figure 7F). In this way, the periodic organization of Ca^2+^ signaling machinery would enable excitatory signals from single spines to be sensed throughout the dendrite, much like the amplifying role of a repeater in telegraph communication.

The regularly arranged distribution of ER-PM junctions with nanometer-range Ca^2+^ sensing/effector capacity shown in this study occurred along all dendritic segments down to the cell body. Previous work from the Blackstone’s^15^ and the Voeltz’s^37^ labs described the presence of a ladder-like ER organization also in axons. The presence of this ER architecture both *in vitro* and *in vivo*, in dendrites and in axons, suggests the existence of neuron-intrinsic mechanisms regulating the formation of this ER architecture in neuronal compartments. Moreover, our results in JPH3-knockdown neurons suggest that voltage-dependent Ca^2+^ channels act as clients rather than determinants at sites where ER protrusions contact the PM.

The laddered-like ER organization may represent a general mechanism for ER assembly in tubular cellular extensions, exhibiting cell- and compartment-specific features whose organizing machinery deserves further investigation. This arrangement suggests a functional mechanism for long-distance intracellular information transfer both in dendrites and axons. In exploring this hypothesis in the future, it will be necessary to further characterize the molecular machinery at JPH3-populated ER-PM junctions, including the roles of other ER-PM tethers, the regulation of STIM1 and STIM2 and their PM partners, and pCaMKII binding sites and molecular targets such as Ca_V_1.2^72–74^. How ER-PM junctions perceive and structurally adapt to changes in PM excitability, for example by altering their morphology, lipid transfer capacity or protein content, will also be an important factor to consider. Finally, it will be worthwhile to determine potential involvement of other organelles like mitochondria, which are often seen closely associated with PM in dendrites, modulate Ca^2+^ signaling at ER-PM junctions through lipid and/or Ca^2+^ exchange pathways.

## Acknowledgments

This work is funded by the Howard Hughes Medical Institute through Janelia Research Campus and the visitor scientist program to L.B. We thank the CellMap Project Team for supporting the volume electron microscopy segmentation. We thank L. Lavis and Janelia Open Chemistry, Tim Brown and the Tool Translation team for generously providing Janelia Fluor dyes reagents. We thank Hyun Ah Yi and Viral Tools; Kevin McGowan and Molecular Genomics; Lihua Wang, Shihing Max Gao and Quantitative Genomics; Jim Cox, Kathy Schaefer, Deepika Walpita, Sabrina Perez, Animal and Biological Services and Kerry Sobieski for providing critical technical support. We would like to thank Kurt Beam for the generous gift of mCherry-JPH3, EYFP-Ca_V_2.1 and GFP-Ca_V_2.2. pORANGE Tubb3-GFP KI was a gift from Harold MacGillavry. pAAV Syn intron Psam4GlyR m3m4 HA was a gift from Scott Sternson. mEmerald-ER-3 was a gift from Michael Davidson. pEX-CMV-SP-YFP-STM2(15-746) was a gift from Tobias Meyer, pmScarlet-I_peroxisome_C1 was a gift from Dorus Gadella. We want to thank C. Hanus for the PSD95-mCherry plasmid and H. Bito for the RCaMP1.07 plasmid. We thank the members of the Lippincott-Schwartz, Ryan and Rangaraju labs for helpful discussions and Carolyn Ott for comments on the manuscript.

## Author contributions

L.B., T.A.R. and J.L.-S. conceived this work and designed the experiments. P.D.C and D.E.C provided critical reagents, advice on experimental protocols, and critical discussion during the development of the project. L.B. designed and performed imaging, pharmacological, biochemical and molecular biology experiments and analyzed data. R.F and V.R. designed and performed functional ER calcium imaging and analysis using two-photon glutamate uncaging experiments. A.S.M performed imaging experiments and data analysis. P.R.H. designed experimental paradigms and conducted RNA extraction, RT-PCR, and data analysis. M.K. generated Julia Pluto.ij notebooks and analyzed data. C.S.X., S.P, and H.F.S. generated FIB-SEM data sets. S.S identified laddered ER in *Drosophila* hemibrain data set, selected and registered the volume for organelle segmentation. A.V.W, G.P, and A.P. performed manual organelles annotations, and P.M.H. produced the video of FIB-SEM rendering. L.B., J.L.-S. and T.A.R wrote the manuscript with input from all co-authors.

## Declaration of interests

The authors declare no competing interests.

## Figures and Figure Legends

**Supplementary Figure 1:**
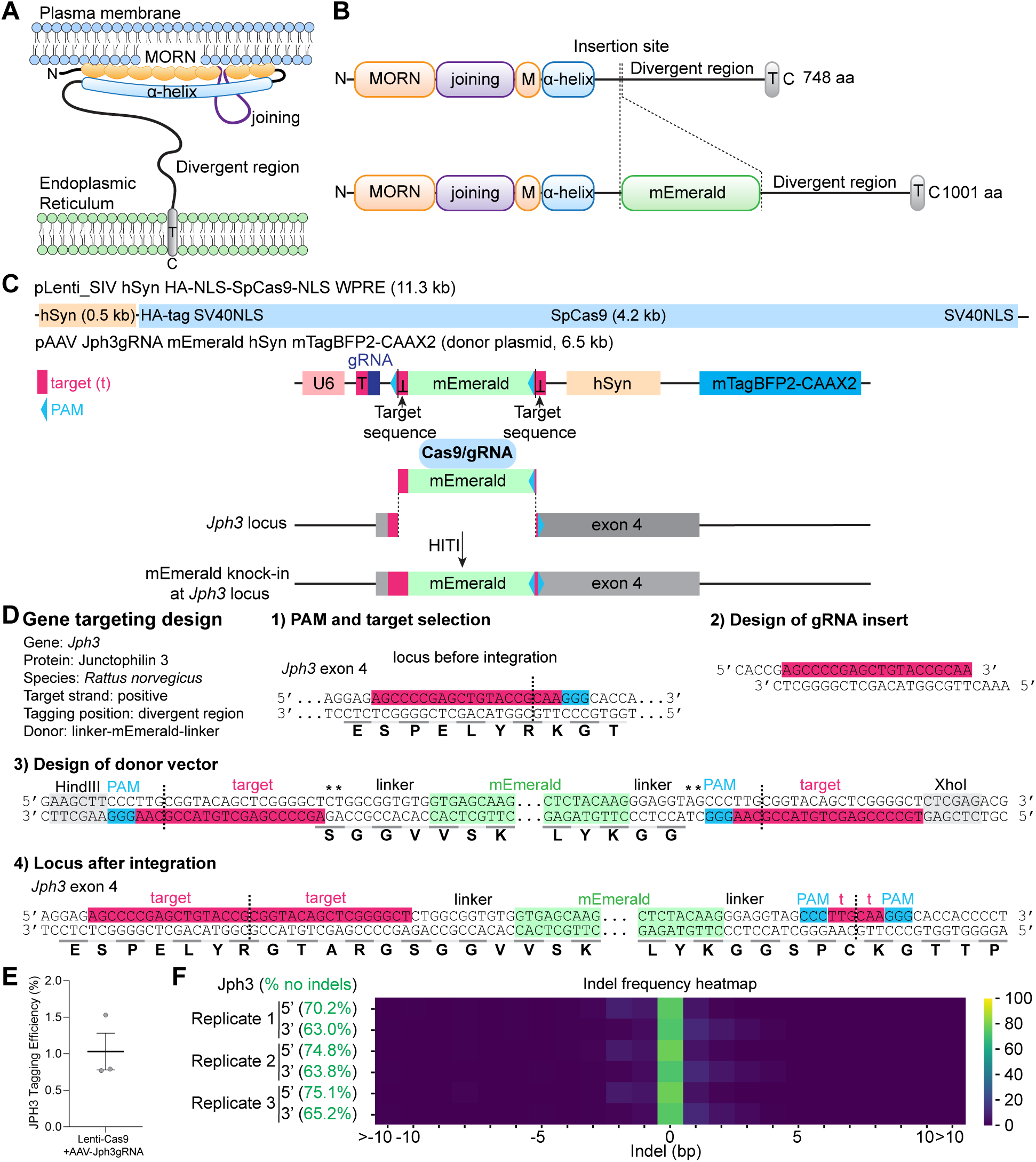
Endogenous tagging of JPH3. A) The N-terminal domain of JPH3 is predicted to contain 8 membrane occupation recognition nexus (MORN) repeats (orange) interrupted by a joining region (purple), capable of interacting with the inner surface of the plasma membrane. Following this domain, an α-helical rich domain (blue) is succeeded by a divergent region characterized by low sequence complexity and predicted to be intrinsically disordered. Finally, a short C-terminal segment traverses the ER membrane (T, gray). This domain organization suggests that JPH3 could be an ER-PM tethering protein. B) Primary sequence organization of JPH3 showing the 8 MORN repeats (orange) interrupted by a joining region (purple), a pseudo-MORN repeat (orange) preceding the α-helical–rich stretch (blue), followed by the divergent region where the RNA guide targets the insertion of mEmerald (green) ending in a short transmembrane helix (T, gray). C) HITI-mediated gene targeting consists of a unique 20-nucleotide RNA guide that directs SpCas9 to the genomic locus of interest. Simultaneously, a donor sequence housing the knock-in gene (mEmerald) is poised for insertion into the targeted genomic locus. In our experiments, we co-infected primary rat neurons using two viral vectors. The first, a lentiviral vector (pLenti_SIV hSyn HA-NLS-SpCas9-NLS WPRE), expressed SpCas9 under control of the human Synapsin 1 promoter. The second vector, an AAV vector (pAAV Jph3gRNA mEmerald hSyn mTagBFP2-CAAX2), included all the elements required for targeted CRISPR/Cas9-based genome editing: a U6-driven expression cassette for the 20 bp guide RNA (gRNA) targeting exon 4 of the rat Jph3 gene, the donor sequence containing the tag of interest (mEmerald) flanked by genomic target sequence homology arms. The gRNA serves a dual purpose: it triggers a genomic double-strand break (DSB) and facilitates the removal of donor DNA from the plasmid, enabling its integration into the genome. Importantly, the orientation of the target sequence and protospacer adjacent motif (PAM) sites flanking the donor is inverted as compared to the genomic sequence guaranteeing that integration occurs in the correct orientation. Additionally, this AAV vector included a plasma membrane marker (mTagBFP2-CAAX2) regulated by the human Synapsin 1 promoter. D) Knock-in construct design for rat Jph3 and endogenous locus sequence after integration. The gRNA targets the sequence in the positive genomic strand of Jph3 exon 4. The target sequence is indicated in magenta, the protospacer adjacent motif (PAM) sequence is in blue. Amino acid translation is shown under the sequences. Dashed lines indicate position of Cas9 cleavage and sites of integration. Asterisks indicate nucleotide added to preserve the open reading frame. E) The endogenous tagging efficiency of our co-infection system was estimated by calculating the ratio between JPH3-mEmerald positive cells and mTagBFP2-CAAX2 expressing neurons (JPH3-mEmerald+/ mTagBFP2-CAAX+) at 21-23 DIV to be 1.03 ± 0.25% (mean ± SEM, N = 3). F) To ensure precise integration of the mEmerald tag into the targeted genomic locus, we conducted next-generation sequencing (NGS) analysis. Our findings indicate successful tagging of JPH3 revealed by a high frequency of in-frame integration in the targeted locus as shown by the heatmap summarizing the sequencing results for 5’ (73.4 ± 1.6%) and 3’ (68.3 ± 0.7%) junction amplicons obtained in 3 independent replicates (mean ± SEM). With correct integration, we observed short out-of-frame insertions or deletions (indels) causing the expression of cytosolic non-fluorescent fragments of the protein or soluble fluorescent fragments due to the loss of the C-terminal transmembrane domain. These observations align with the rarity of off-target integration observed, which mainly occurred as soluble mEmerald expression. Heatmap is color-coded for the frequency of indel size, as analyzed using CRIS.py.

**Supplementary Figure 2:**
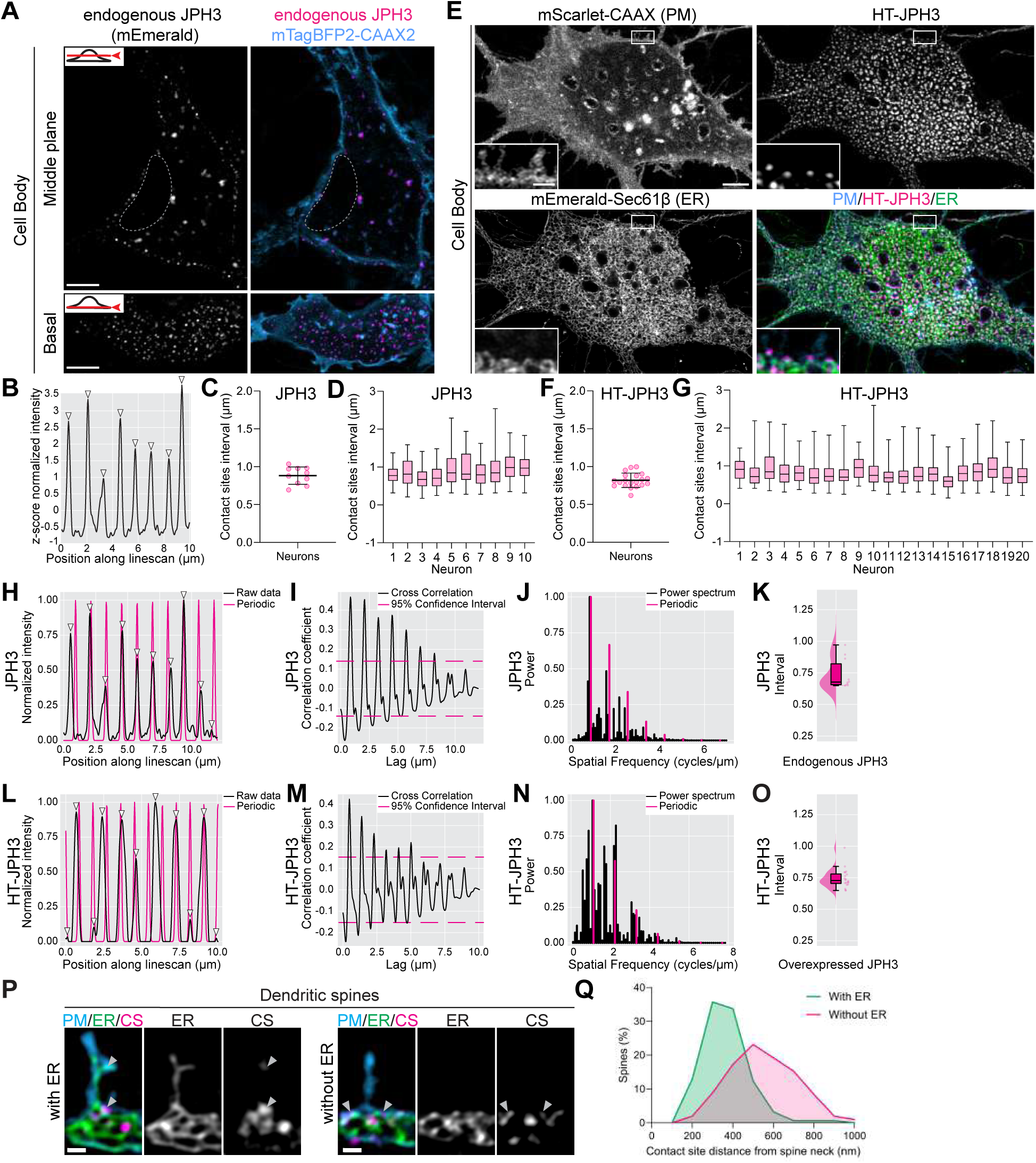
Junctophilin-3 accumulates at quasi-periodically arranged ER-PM contacts in dendrites of primary rat hippocampal neurons. A) JPH3 endogenously tagged with mEmerald in the cell body of a primary hippocampal neuron. Endogenous JPH3 accumulates in puncta juxtaposed to the plasma membrane (mTagBFP2-CAAX2). Scale bar: 5 μm. B) Normalized intensity plot profile of endogenously tagged JPH3 hotspots drawn along the dendritic plasma membrane. Arrows indicate well-separated, evenly spaced peaks detected based on an unbiased minimum prominence threshold. C) JPH3 puncta are localized at evenly spaced intervals of ∼1 μm (0.88 ± 0.12 μm, n = 10 neurons, N = 5 animals, mean ± SD) along the entire dendritic arbor. D) Box plot showing endogenous JPH3 contact sites intervals regularly spaced at ∼1 μm in all neurons analyzed (43 to 122 contact intervals were analyzed in each neuron). E) Overexpression of HaloTag(HT)-JPH3 (labeled with JF647 HaloTag-ligand) in primary rat neurons at 21 DIV showed accumulation at ER-PM contact sites similar to endogenously tagged JPH3. Overexpression conditions resulted in a brighter, more consistent JPH3 signal and enlarged ER-PM junctions. Scale bar: 5 μm, inset scale bar: 1 μm. F) Average contact sites interval measured as the distance between adjacent HT-JPH3 clusters in intensity plot profiles drawn along the dendritic plasma membrane, as shown in panel B (0.82 ± 0.10 μm, n = 20 neurons, N = 3 animals, mean ± SD) G) Box plot showing regular HT-JPH3 contact sites spacing of ∼1 μm in all neurons analyzed (56 to 118 contacts were analyzed in each neuron). H,L) Upon detecting peaks in the line scans of both endogenous JPH3 (H) and overexpressed HT-JPH3 (L), we noticed that the intervals between the junctions were spaced at very regular intervals (panels C,D for endogenous JPH3 and F,G for HT-JPH3), suggesting some degree of periodicity. To evaluate the degree of periodicity, we synthesized a localization signal with maxima at the detected peaks and cross-correlated them with periodic functions of comparable frequencies (the periodic Gaussian signal with the matched detected frequency is plotted in magenta in H for endogenous JPH3 and in L for HT-JPH3). We found that the localization signals significantly correlated with the frequency-matched periodic functions when compared to a localization signal generated by a white noise process (as shown in I and M for endogenous and HT-JPH3, respectively, where the dashed magenta lines indicate the 95% confidence intervals as described in the Methods). The normalized Fourier power spectrum of the periodic signal and the synthetic signal with Gaussians placed at the detected peaks is shown in (J) for endogenous and (N) for HT-JPH3. Raincloud plots of the contact sites intervals in dendrites detected from Fourier analysis are shown in (K) for endogenous and (O) for overexpressed JPH3. While not purely periodic in the mathematical sense due to biological variability in the intervals, these results show that the junction membrane complexes marked by JPH3 occur at regular intervals distinct from pure chance illustrating a quasi-periodic distribution. P) Lattice-SIM images illustrate the localization of contact sites (CS), marked by accumulation of HaloTag-JPH3 (labeled with JF635 HaloTag-ligand) relative to dendritic spines (visualized with the plasma membrane (PM) marker mScarlet-CAAX2), with or without ER (mEmerald-Sec61β) accumulation in the spine head. Scale bars: 1 µm. Q) Analysis of contact sites distance from the spine head reveals that when ER tubules protrude in the spine head, contact sites are located approximately 363 ± 116 nm from the spine neck (n = 154 spines). In spines with no ER presence, the closest contact site is localized approximately 566 ± 191 nm from the spine neck (n = 203 spines). Data were analyzed in 10 neurons from 3 independent experiments and represented as mean ± SD.

**Supplementary Figure 3:**
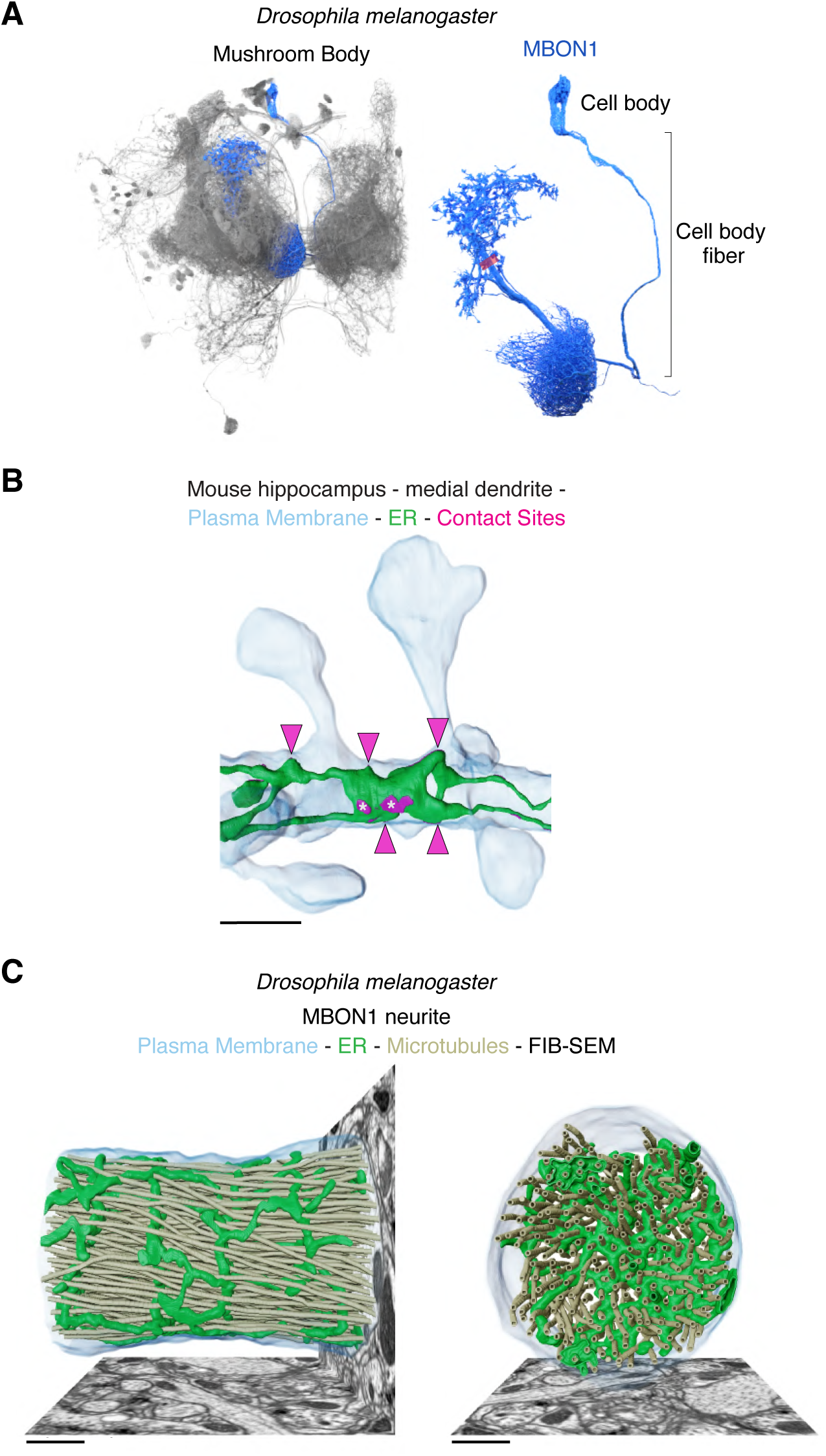
ER-PM junctions stabilize ER arrays in vertebrate and invertebrate neurons. A) Illustration created by overlaying 3D renderings of selected neurons from the FIB-SEM data set encompassing the *Drosophila melanogaster* Mushroom Body (MB). Organelles segmentation shown in Figure 2 and Figure 3 was conducted within the volume highlighted with a magenta box, corresponding to a region of the MBON1 neurite shown in blue whose position in the MB and structural organization is also depicted. B) 3D renderings of ER (green), plasma membrane (blue) and ER-PM contact sites (magenta) in a mouse hippocampal neuron segmented from FIB-SEM datasets. Magenta arrowheads highlight contact sites anchoring ER arrays to the plasma membrane. Asterisks indicate non-rung associated contact sites. Scale bar: 1 μm. C) Longitudinal and orthogonal views of the 3D renderings of ER (green), plasma membrane (blue), and microtubules (tan) in *Drosophila* MBON1 neuron segmented from FIB-SEM datasets. Scale bars: 1 µm.

**Supplementary Figure 4:**
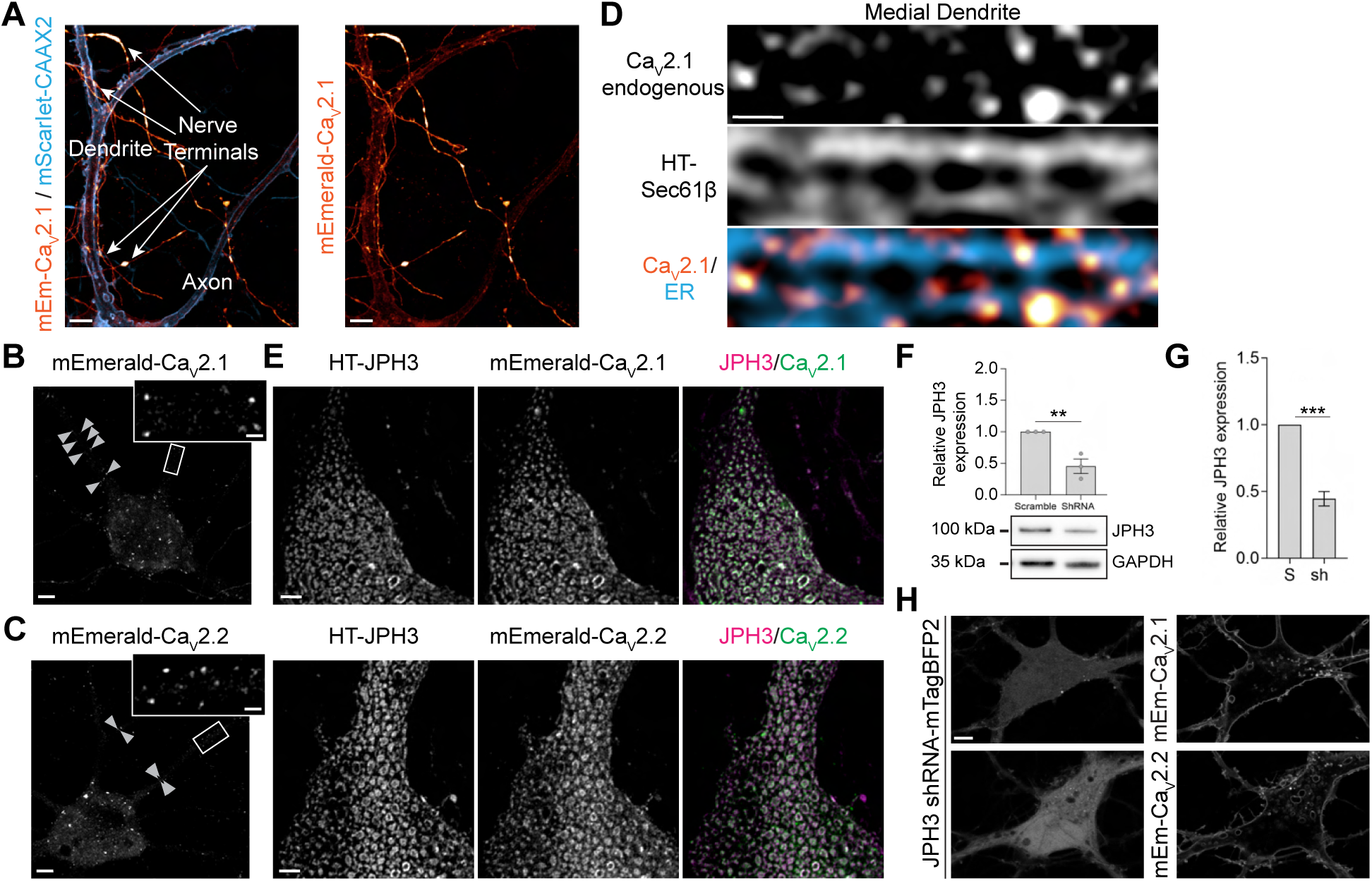
Effects of Junctophilin-3 expression on voltage-gated calcium channels recruitment to ER-PM junctions. A) Live-cell imaging of mEmerald-Ca_V_2.1 (glow LUT) and mScarlet-CAAX2 (cyan LUT) in primary rat hippocampal neurons showed mEmerald-Ca_V_2.1 accumulation in nerve terminals and dendrites. Scale bar: 5 μm. B-C) Both mEmerald-Ca_V_2.1 and mEmerald-Ca_V_2.2 accumulate in small clusters in neuronal cell bodies and dendrites. In large proximal dendrites, Ca_V_2.1 and Ca_V_2.2 clusters are rare, but they are often symmetrically distributed on opposite sides of the dendritic membrane (examples are indicated with arrowheads and shown in insets). Scale bar: 5 μm. D) Immunostaining experiments revealed endogenous Ca_V_2.1 puncta positioned at opposite sides of ER rungs visualized using the ER marker HaloTag-Sec61β (labeled with JF647 HaloTag-ligand). Scale bar: 0.5 μm. E) mEmerald-Ca_V_2.1 and mEmerald-Ca_V_2.2 colocalized with HaloTag-JPH3 (labeled with JF635 HaloTag ligand) in neuronal cell bodies. Scale bar: 5 μm. F) ShRNA knockdown of JPH3 significantly reduced protein levels in Western blot experiments. Relative JPH3 expression measured after knockdown = 0.45 ± 0.11, N = 3. Data analyzed with unpaired t-test G) RT-PCR shows significant downregulation of JPH3 transcripts upon JPH3 shRNA expression. Relative JPH3 expression = 0.45 ± 0.05, n=7, N=3. Data were analyzed with the Mann-Whitney-test. H) Knockdown of JPH3 (neurons expressing shRNA for JPH3 were identified by cytosolic expression of mTagBFP2) results in the loss of mEmerald-Ca_V_2.1 and mEmerald-Ca_V_2.2 clusters in cell bodies. Scale bar: 5 μm. All data are mean ± SEM; ** p<0.01, *** p<0.001.

**Supplementary Figure 5:**
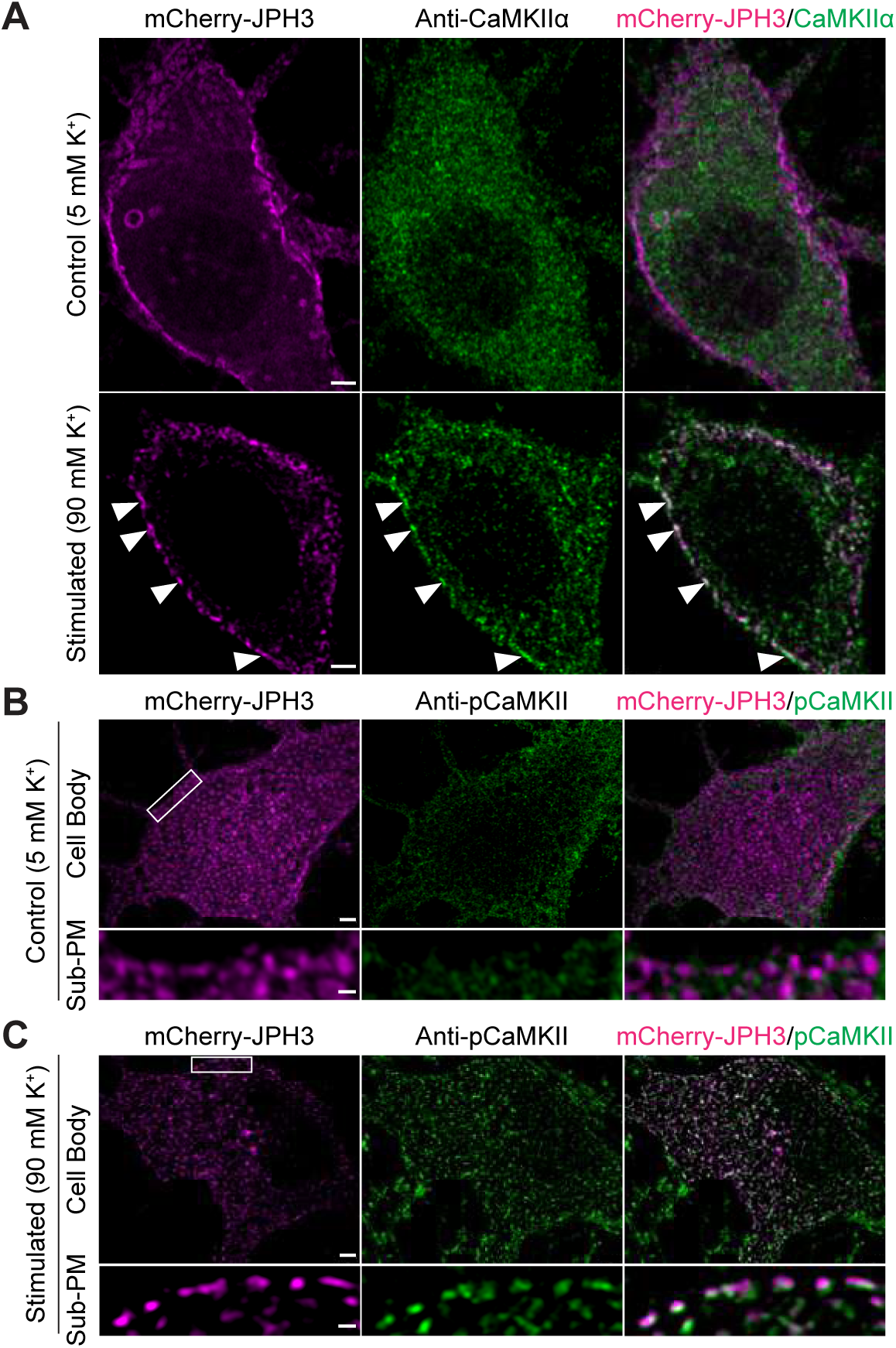
CaMKIIα and pCaMKII accumulate at ER-PM contacts in cell bodies after neuronal stimulation. A) Immunofluorescence assays designed to ascertain the localization of the α isoform of CaMKII before and after depolarization, showed CaMKIIα accumulation at ER-PM contacts (arrowheads) only after neuronal stimulation. Scale bar: 2 μm. B-C) Immunofluorescence analysis of pCaMKII distribution before (B) and after stimulation with 90mM K^+^ (C), reveals pCaMKII hotspots of accumulation at mCherry-JPH3 junctions juxtaposed to the PM (Sub-PM) within the neuronal cell body. Scale bar: 2 μm main panels, 0.5 μm insets.

**Supplementary Figure 6:**
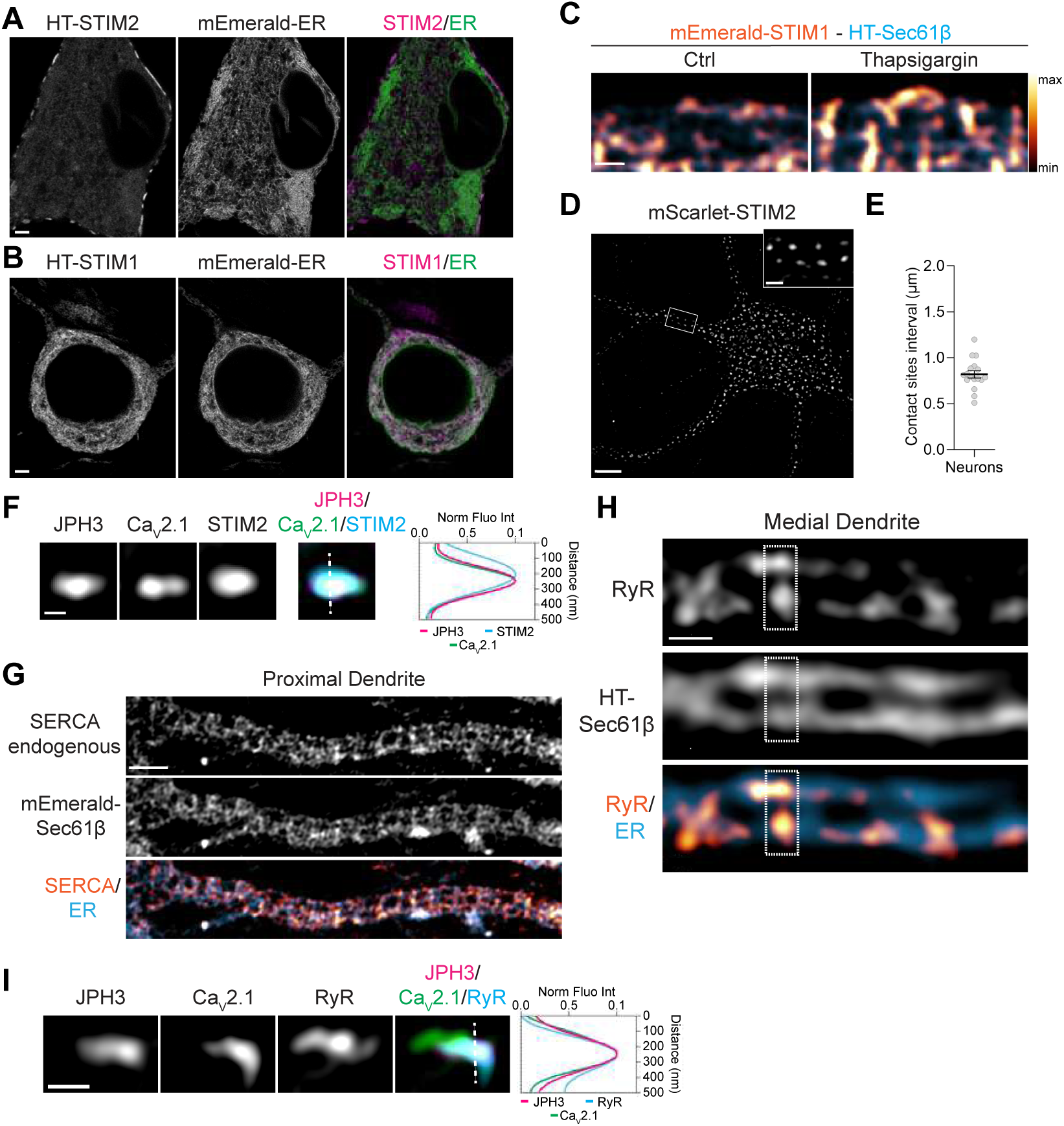
Molecular machinery for the fine-tuning of ER Ca^2+^ release and uptake populate regular contact sites in neuronal dendrites. A-B) Cell bodies of primary neurons at 21DIV co-expressing HaloTag-STIM2 (A) and HaloTag-STIM1 (B) with the ER marker mEmerald-ER. While HaloTag-STIM2 accumulated at ER-PM contacts, HaloTag-STIM1 distributed in the ER meshwork in basal conditions. Scale bar: 0.2 μm. C) HaloTag-STIM1redistributed at ER-PM contacts upon ER Ca^2+^ depletion with thapsigargin. Scale bar: 1 μm. D) mScarlet-STIM2distributes at evenly spaced contacts in neuronal dendrites. Scale bar: 5 μm main panel, 1 μm inset. E) Average contact sites interval measured as the distance between adjacent mScarlet-STIM2clusters along the dendritic plasma membrane (0.82 ± 0.16 μm, n = 17, N = 3). F) mCherry-JPH3 (JPH3), mEmerald-Ca_V_2.1 (Ca_V_2.1) and Halo-STIM2 (STIM2) colocalize at periodic ER-PM contacts in dendrites. Scale bar: 0.2 μm. G) Immunofluorescence experiments revealed that endoplasmic reticulum Ca^2+^ ATPase (SERCA) pump uniformly distributes in dendritic ER. Scale bar: 2 μm. H) Immunolabeling experiments revealed that RyRs localized at the extremities of ER rungs. The dashed box highlights a frequent correlation between RyRs hotspots’ fluorescence intensity at opposite junctional sites of ER rungs. Scale bar: 0.5 μm. I) Immunofluorescence experiments revealed the colocalization of RyRs, HaloTag-JPH3 _(_JPH3) a_n_d mEmerald-Ca_V_2.1 (Ca_V_2.1) at periodic ER-PM contacts in dendrites. Scale bar: 0.2 μm. HaloTag was labeled with JF635 or JF647.

**Supplementary Figure 7:**
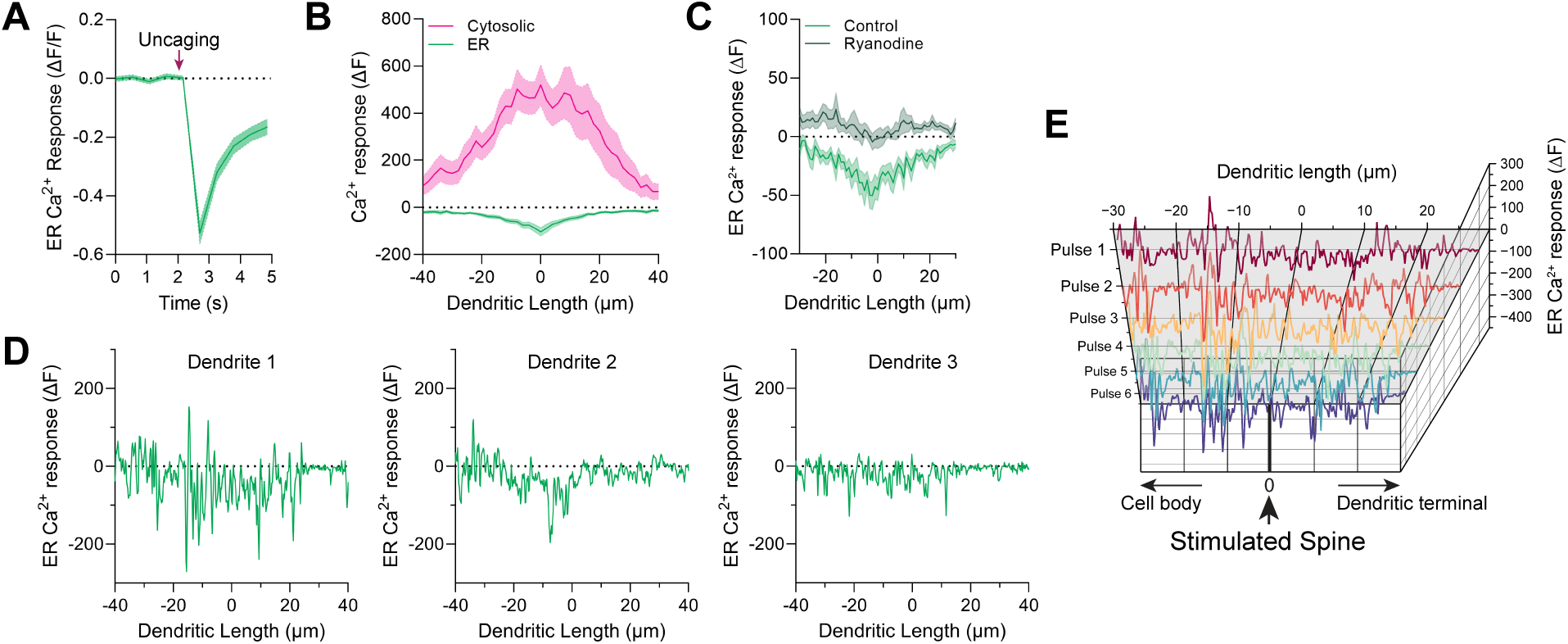
Regular ER-PM contacts are sites of Ryanodine-dependent ER Ca^2+^ release. A) Average temporal profile measured for the initial ER Ca^2+^ release event detected in each dendrite shown as mean ± SEM (n = 19 dendrites, N = 3 biological replicates). B) Spatial profile of cytosolic Ca^2+^ measured using RCaMP1.07 (magenta) showed an elevation in cytosolic Ca^2+^ concurrent with Ca^2+^ release from dendritic ER detected using the ER Ca^2+^ indicator ER-GCaMP6-210 (green) upon local spine stimulation. Sample size for cytosolic Ca^2+^ profile: n = 24 dendrites, N = 2 biological replicates; sample size for luminal ER Ca^2+^ profile: n = 31 dendrites, N = 3 biological replicates. Data represented as mean ± SEM. C) Average dendritic ER Ca^2+^ profile measured during local spine stimulation in control and after ryanodine treatment (n=14 dendrites, N = 3 biological replicates). Data represented as mean ± SEM. D) Representative examples of ER Ca^2+^ spatial profiles measured in three individual dendrites illustrating ER Ca^2+^ dynamics. E) Sequential spatial profiles resulting from 6 consecutive uncaging events within the same dendrite.

## Video Legends

**Video 1**: Z-stack of primary rat hippocampal neuron (21 DIV) expressing JPH3 endogenously tagged with mEmerald. JPH3 is distributed in small puncta juxtaposed to the plasma membrane. Scale bar: 2 μm. Z-step size: 0.133 μm.

**Video 2**: 3D renderings of the endoplasmic reticulum (ER - green), plasma membrane (blue), mitochondria (pink), microtubules (tan), and ER-plasma membrane contacts (magenta) segmented from FIB-SEM datasets of a *Drosophila melanogaster* MBON1 neuron.

**Video 3**: Time-lapse acquired using 2D lattice-SIM in burst mode of HaloTag-Sec61β (labeled with JF585 HaloTag-ligand) expressing neurons, revealing the dynamic nature of ER tubules in contrast with the persistence of ER-plasma membrane junctional sites over time. Scale bars: 0.5 μm.

**Video 4**: Time-lapse acquired using Lattice-SIM of mEmerald-Sec61β expressing neurons. Lysosomes were stained using Lysview650. Lysosomes swiftly move through ladder-like ER arrays. Scale bars: 0.5 μm.

## Materials and methods

### Ethics statement

All experimental procedures involving the use of rats were performed in compliance with the *Guide for the Care and Use of Laboratory Animals*. All experiments were approved by the Janelia Research Campus (JRC) and Max Planck Florida Institute (MPFI) Institutional Animal Care and Use Committee (IACUC) and Institutional Biosafety Committee (protocol number 21-206). JRC is an AAALAC-accredited institution. Timed-pregnant rats (Sprague Dawley) used in the experiments were obtained from Charles River Laboratories (Wilmington, MA) and maintained under specific pathogen-free conditions.

### Neuronal Cultures

Primary cultures of hippocampal neurons were prepared from rat brains of both genders at embryonic day 19 (E19). The hippocampi were dissected in cold Hank’s Balanced Salt Solution (Sigma, H2387), which was supplemented with 1 mM HEPES (Fisher, BP310), 4 mM NaHCO_3_ (Sigma-Aldrich S5761), and 20% FBS (Biotechne, S11550H). The cells were then dissociated by tissue trituration and papain treatment (Worthington Biochemical Corporation, LK003178, PDS Kit) in HBSS at pH 7.4 at 37°C for 10 minutes. Live, dissociated cells (Trypan Blue exclusion) were counted and plated on 25 mm round coverslips (Electron Microscopy Science, 72196-25) coated with Poly-L-Ornithine Solution (0.01%, Sigma-Aldrich, P4957) and 10 μg/ml laminin (Sigma-Aldrich, 11243217001) in DPBS (Thermofisher, 14040182) at a density of 3.4 x 10^4^ cells/cm^2^ in NeuroCult^TM^ Neuronal Plating Medium (STEMCELL Technologies, 05713) supplemented with 2% NeuroCult^TM^ SM1 neuronal supplement (STEMCELL Technologies, 05711) and 2 mM GlutaMAX™ Supplement (Gibco, 35050061). Five days after plating, half of the plating medium was removed from each well and replenished with the same volume of BrainPhys™ Neuronal Medium (STEMCELL Technologies, 05792) supplemented with 2% NeuroCult™ SM1 Neuronal supplement. The cells were maintained *in vitro* at 37°C and 5% CO_2_. From DIV 5 onwards, the medium was refreshed every four days by replacing half of the medium with BrainPhys™ neuronal medium supplemented with 2% NeuroCult™ SM1 neuronal supplement.

### Plasmids

pAAV hSyn INTRON mEmerald-Sec61β WPRE, pAAV hSyn INTRON HaloTag-Sec61β WPRE, pAAV hSyn INTRON mEmerald-ER WPRE, pAAV hSyn INTRON mTagBFP2-CAAX2 WPRE, pAAV hSyn INTRON mScarlet-CAAX2 WPRE, pAAV hSyn INTRON HaloTag-STIM1 were generated subcloning the cORF of mEmerald-Sec61β (from Addgene #90992), HaloTag-Sec61β (Addgene #123285), mEmerald-ER3 (Addgene #54082), mTagBFP2-CAAX2 (synthesized with gBlock), mScarletI-CAAX2 (synthesized with gBlock), HaloTag-STIM1 (synthesized with gBlock) at BamHI and HindIII sites of pAAV Syn intron Psam4GlyR m3m4 HA (a gift from Scott Sternson, Addgene #196041). pAAV hSyn INTRON mEmerald-STIM1 was generated subcloning SS-mEmerald-linker (synthesized with gBlock) at BamHI and BsrGI restriction enzyme sites of pAAV hSyn INTRON HaloTag-STIM1. STIM2 construct design was based on SP-YFP-STM2(15-746) (Addgene #18862). We replaced YFP with HaloTag and mScarletI (from Addgene #85065) and subcloned the full cORF in AAV backbone at SpeI and HindIII sites to generate pAAV hSyn INTRON HaloTag-STIM2 WPRE and pAAV hSyn INTRON mScarlet-STIM2 WPRE. pORANGE Tubb3-GFP KI was obtained from Addgene (Addgene plasmid #131497). The AAV vector pAAV hSyn INTRON mCherry-Jph3 WPRE, expressing human Jph3 gene fused to the fluorescent protein mCherry, was generated by subcloning the cORF of mCherry-JPH3, a kind gift from Kurt Beam (University of Colorado School of Medicine, CO, USA), into our AAV template backbone (Addgene #196041) at BamHI and HindIII cloning sites. pAAV hSyn INTRON HaloTag-Jph3 WPRE was generated replacing mCherry with HaloTag (from Addgene #123285) at BglII and HindIII sites. pLenti_SIV-(SalI)-syn-IRES-mEmerald-Ca_V_2.1 WPRE and pLenti_SIV-(SalI)-syn-IRES-mEmerald-Ca_V_2.2 WPRE were generated cloning Ca_V_2.1 and Ca_V_2.2 cORF fused to mEmerald at the N-terminus in a third-generation, simian immunodeficiency based, pro**-**lentiviral vector^1^ at AgeI and HpaI sites. EYFP-Ca_V_2.1 and GFP-Ca_V_2.2 were kind gifts of Kurt Beam (University of Colorado School of Medicine, CO, USA). ER-GCaMP6-210 is described in de Juan-Sanz et al., 2017^2^ (Addgene #86919). The post-synaptic marker PSD95-mCherry is described in Hanus et al., eLife 2016^3^. RCaMP 1.07 was obtained from Bito H and described in Inoue et al., Nature Methods 2014^4^. For all these clones, PCR amplification of the fragments and their subsequent ligation was performed using the In-Fusion cloning kit and online tools (BD Clontech, Takara Bio, USA), or they were synthesized by GenScript (Piscataway, NJ, USA). All plasmids were verified by sequencing by GenScript or Plasmidsaurus, (Eugene, OR, USA).

### Adeno-associated virus (AAV) and lentivirus particle production and applications

The Janelia Viral Tools team at HHMI Janelia Research Campus (Ashburn, VA) produced adeno-associated virus (AAV) and lentivirus particles. To generate pLenti-SIV, a VSV-G envelope-expressing plasmid (pMD2.G, Addgene 12259) and SIV_Gag_Pol and SIV_Rev plasmids (these two constructs will be available in Addgene upon publication of this manuscript) were co-transfected. AAV or lentivirus particles were added to DIV3 or DIV5 neurons, and media exchange was performed 2-4 days after infection. All experiments were conducted after 13 days of viral transduction.

### Lipofectamine transfection

Primary neurons were transfected at 5 days in vitro (DIV) using Lipofectamine 2000 Transfection Reagent (Invitrogen, 11668027) following the manufacturer’s instructions. The neuronal conditioned media was first collected and replaced with warm Neurobasal™ medium (ThermoFisher Scientific, 21103049). A transfection mix was prepared using a 1:2 DNA to Lipofectamine ratio in Neurobasal™ medium, which was added dropwise to the cells. The cells were then incubated with the transfection mix for 30 minutes. Subsequently, the transfection medium was replaced with neuronal-conditioned media, and the cells were maintained *in vitro* at 37°C and 5% CO_2_.

### Design and generation of viral vectors for JPH3 endogenous tagging

We used the homology-independent targeted integration (HITI) method to fluorescently label JPH3 leveraging on non-homologous end joining (NHEJ) for endogenous protein tagging in postmitotic neurons^5^. We cloned a third-generation, simian immunodeficiency-based, pro-lentiviral vector^1^ containing a 476 bp human synapsin-1 promoter element driving the neuronal specific expression of HA-tagged SpCas9 flanked by the bipartite SV40 nuclear localization signal (NLS) HA-NLS-SpCas9-NLS (from Addgene, plasmid #131497) at AgeI and BsrGI sites followed by woodchuck hepatitis post-transcriptional regulatory element (WPRE) inserted between the pro-lentiviral long terminal repeats (pLenti_SIV_hSyn_HA-NLS-SpCas9-NLS WPRE). We synthesized an AAV knock-in template vector (pAAV KI Cloning Vector_hSyn mTagBFP2-CAAX2) using pAAV hSyn intron Psam4GlyR m3m4 HA (Addgene, plasmid #196041) as backbone containing: 1) a U6-driven expression cassette for the guide RNA (gRNA) targeting the genomic locus of interest that can be inserted at BbsI sites; 2) HindIII and XhoI sites where it is possible to subclone the donor sequence containing the (fluorescent) tag (the template for this region was the Addgene plasmid #131497, pORANGE Tubb3-GFP KI); 3) a 448 bp human synapsin-1 promoter element driving the neuronal specific expression of 4) the fluorescent protein mTagBFP2 (Addgene plasmid #191566 pLKO.1-TRC mTagBFP2) fused to the plasma membrane targeting sequence CAAX2 (Addgene plasmid #162247, eMagAF-EGFP-PM). This construct was synthesized by GenScript. We decided to introduce the mEmerald fluorescent protein tag after the N-terminal MORN repeats and α-helix domain of JPH3^6^ specifically targeting the divergent region of the protein, a region of low sequence complexity predicted to be intrinsically disordered (Figure S1A and S1B). 20 nt RNA guides were designed and analyzed using the CRISPR tool of Benchling (Benchling, Biology Software, 2024, retrieved from https://benchling.com) in the exon 4 of rat Jph3 genomic sequence (mRatBN7.2) downloaded from useast.ensembl.org (Ensembl version: ENSRNOG00000018784.7) using Rnor_6.0_EnsemblRelease_104 (Rattus norvegicus) as reference genome and NGG (SpCas9, 3’side) PAM. The sgRNA guide was chosen, taking into consideration on- and off-target optimized scores from Doench, Fusi et al. (2016)^7^. The 20-bp gRNA selected (AGCCCCGAGCTGTACCGCAA) was inserted into AAV knock-in template vector at BbsI sites (Figure S1C and S1D). The donor sequence was designed to contain the sequence of the fluorescent protein tag mEmerald flanked by two Cas9 target sites identical to the genomic target site and inserted at HindIII and XhoI sites. Importantly, to facilitate genomic integration of the donor sequence in the correct orientation, these target sites, including PAM sequences, were inserted as the reverse complement of the genomic target sequence (Figure S1C and S1D). Short linker sequences of at least three amino acids and additional base pairs to make the donor in frame after integration in the genome were introduced between the target sites and the tag sequence (Figure S1C and S1D). At 3 days in vitro (DIV), primary rat neurons were coinfected with the lentiviral SpCas9 vector and AAV1 vector carrying the RNA guide for the rat Jph3 gene targeting, the donor tag, and a plasma membrane marker (mTagBFP2-CAAX2). This allowed the visualization of the plasma membrane of all neurons expressing the gRNA and the labeling of JPH3 in a subset of neurons at 19-23 days in vitro (DIV). The efficiency of JPH3 tagging at endogenous locus was measured in primary neuronal cultures at 21 DIV. Cells were fixed and immunolabeled with anti-GFP primary Ab (Aves, GFP-1020) and AlexaFluor^TM^ 594 goat-anti-chicken secondary antibody (Invitrogen, A11042). Fields of view tiling almost the entire coverslip were acquired with a BZ-X810 Keyence system (Keyence corporation of America, Itasca, IL) with a Plan Apochromat 10x/0.45 (BZ-PA10) objective, LED illumination and CCD camera. mTagBFP2-CAAX2 fluorescent signal was detected with BZ-X filter DAPI (OP-87762, excitation wavelength 360/40, emission wavelength 460/50, dichroic mirror wavelength 400). AlexaFluor^TM^ 594-secondary antibody was detected with BZ-X filter TexasRed (OP-87765, excitation wavelength 560/40, emission wavelength 630/75, dichroic mirror wavelength 585). Endogenous tagging efficiency was measured as the ratio between the number of cells expressing JPH3-mEmerald and all the mTagBFP2-CAAX2 positive cells.

### Next Generation Sequencing (NGS) of gRNA genomic sites of integration

Genomic DNA was isolated from neurons at 21 DIV, after coinfection with pLenti_SIV_hSyn_HA-NLS-SpCas9-NLS_WPRE and pAAV_Jph3gRNA_mEmerald_hSyn_mTagBFP2-CAAX2 at 3 DIV. DNeasy Blood & Tissue Kit (Qiagen, 69504) was used for DNA extraction following the manufacturer’s instructions. Genomic PCR was performed to amplify the integrated transgene’s 5′ and 3′ junctions. PCR primers were designed and selected by using NCBI primer-BLAST with an insertion size of approximately 200 bp. The forward and reverse primers were appended with the Illumina P5 and P7 adaptor sequences, respectively. Additionally, 8 bp index barcodes were introduced at the P7 end for sample identification. To maintain sequence diversity, a mix of 10 P5-primers with stagger regions of different length was designed. All primers were ordered from Integrated DNA Technologies (IDT). The PCR was carried out using a high-fidelity amplification kit, NEBNext Ultra II Q5 Master Mix (New England Biolabs, M0544). The resulting PCR products were purified through double-sided AMPure XP beads cleanup (Beckman Coulter, A63882), followed by bioanalyzer quality control to ensure the obtainment of clean PCR products with appropriate size. The amplicons were pooled at an average concentration of 1 nM and subjected to sequencing on Illumina Miniseq platform using single-end 200 bp reads. Sequencing results were analyzed using CRIS.py^8^. Indel frequencies were plotted in a heatmap as the average percentage from the forward and reverse reads. The number of forward and reverse reads were averaged per junction for each knock-in and plotted. Average number of reads obtained with deep sequencing for 5’ junction: 1.52 × 10^6^ reads ± 0.15 × 10^6^; 3’ junction: 7.5 × 10^5^ ± 0.32 × 10^5^, mean + SD).

### Live-cell imaging experiments

Live imaging experiments were conducted in two different media: BrainPhys™ Imaging Optimized Medium (STEMCELL Technologies, 05796) supplemented with 2% NeuroCult™ SM1 neuronal supplement (STEMCELL Technologies) at a temperature of 37 °C and 5% CO_2_, or Tyrode’s solution, which contained 150 mM NaCl, 5 mM KCl, 2 mM CaCl_2_, 2 mM MgCl_2_, 10 mM HEPES and 10 mM glucose, and was buffered to pH 7.4. The experiments were performed using Elyra 7 (Zeiss) or Zeiss LSM 980 with Airyscan (Zeiss).

### HaloTag labeling

To label neurons with HaloTag fusion constructs, cells were incubated with 50-100 nM JF HaloTag-ligand in complete BrainPhys™ Imaging Optimized Medium supplemented with 2% NeuroCult™ SM1 neuronal supplement for 30 minutes at 37 °C and 5% CO_2_. After incubation, neurons were rinsed three times and incubated in complete BrainPhys™ Imaging Optimized Medium for a minimum of 30 minutes at 37°C and 5% CO_2_ to remove the dye. Subsequently, the cells were rinsed again and prepared for imaging.

### Lattice-Structured Illumination Microscopy (Lattice-SIM)

All experiments utilizing Lattice-SIM were conducted using the Elyra 7 KMAT microscope system from Zeiss. The microscope was built around an inverted AxioObserver 7 controlled using ZEN 3.0 SR software. Imaging was carried out at 37°C and 5% CO_2_ using an alpha Plan-Apochromat 63x/1.46 Oil Korr M27 Var2 objective. A 488 nm laser was used to excite mEmerald, a 561 nm laser was used for mScarlet or JF585 HaloTag-ligand, and a 642 nm laser was used for JF635 or JF646 HaloTag-ligands and SirActin (SiR-Actin Kit, Cytoskeleton Inc, CY-SC001). Fluorescence emission was detected by pco.edge CLHS sCMOS cameras with appropriate bandpass filters LBF 405/488/561/642 + SBS LP 560 nm, BP 495-550 + 570-620 nm + SBS LP 560 and BP 420-480 + BP 495-525 + LP 655 nm + SBS LP 560, respectively. Fields of view of 123.9 x 123.9 μm (1280 x 1280 pixels) at 0.133 μm z-steps with 13 phase images were acquired. 3D channel alignment was performed on high-resolution z-stacks of TetraSpeck^TM^ Microspheres (ThermoFisher; T14792) imaged with the same parameters. Lattice SIM processing was performed using ZEN 3.0 SR software with 3D default parameters.

Rapid dynamics of the ladder-like ER in medial dendrites were generated from 2D-time series of neurons expressing HaloTag-Sec61β labeled with JF585 and imaged with a 561 nm laser and BP 495-550 + 570-620 nm + SBS LP 560 bandpass filters. Time series with no interval delay were acquired and processed with 2D Lattice SIM default parameters and Burst Mode, generating time series of 20 ms intervals.

Lysosomes and ER dynamics were imaged in neurons expressing mEmerald-Sec61β and incubated for 30 minutes with LysoView^TM^ 650 (Biotium, 70059) in imaging medium. 2D Lattice-SIM time series were acquired simultaneously detecting mEmerald and LysoView^TM^ 650 emission with LBF 405/488/561/642 + SBS LP 560 upon 488 nm and 642 nm laser excitation. Fields of view of 49.6 x 49.6 μm (512 x 512 pixels) were imaged at 60 Hz for 5 minutes.

Neurons transiently transfected with pORANGE Tubb3-GFP KI at DIV 3 and subsequently infected with HaloTag-Sec61β (labeled with JF585-HaloTag-ligand) were imaged at 21 DIV. Nocodazole-treated cells were incubated with 5 mM nocodazole (ThermoFisher Scientific, AC358240100) for 2-3 h at 37°C and 5% CO_2_ before imaging. Endogenous labeling for β3-tubulin was used to assess the extent of depolarization of the microtubule network in the cells imaged. 3D Lattice-SIM images were acquired sequentially detecting JF585 and mEmerald emission with BP 495-550 + BP 570-620 + SBS LP 560 and LBF 405/488/561/642 + SBS LP 560 upon 488 nm and 561 nm laser excitation. Fields of view of 80.1 x 80.1 μm (1280 x 1280 pixels) at 0.110 μm z-steps with 9 phase images were acquired.

### Airyscan microscopy

Airyscan experiments were conducted on an inverted Zeiss LSM 980 microscope equipped with an Airyscan module. To excite different fluorescent proteins/dyes, lasers emitting 639 nm, 561 nm, 488 nm and 405 nm excitation wavelengths were used. More specifically, JF646 or JF635 were excited using the 639 nm laser; JF585, mCherry, mScarlet or AlexaFluor^TM^ 594 using the 561 nm laser; mEmerald or AlexaFluor^TM^ 488 using the 488 nm laser; mTagBFP2 using the 405 nm laser.

Imaging of endogenously tagged JPH3 with mEmerald and mTagBFP2-CAAX2 signals were imaged in 18-21 DIV neurons with sequential detection of mEmerald and mTagBFP2 emission with MBS 488/639 and MBS-405 + SBS SP 505, respectively. Z-stacks of 79.2 x 79.2 μm (2245 x 2245 pixels) were imaged.

Neurons expressing mScarlet-CAAX2, mEmerald-Ca_V_2.1 or mEmerald-Ca_V_2.2 and pLKO.1 Scramble shRNA-mTagBFP2 (Control) or pLKO.1 JPH3 shRNA-mTagBFP2 were imaged sequentially with MBS 488/561 + BP 420-480 + BP 495 + 550 and MBS 408 + SBS SP 505. Imaging of Z-stacks of 79.2 x 79.2 μm (2245 x 2245 pixels) was performed. mCherry-JPH3 contact site intervals and size in neuronal dendrites were imaged in neurons expressing mCherry-JPH3, mEmerald-Sec61β and mTagBFP2-CAAX2 at 18-22 DIV following a 12-hour treatment with 0.5 μM TTX (Tetrodotoxin citrate, Abcam, ab120055) to reduce neuronal activity, or with 20 μM Bicuculline (Sigma-Aldrich, 14340) or 250 nM Kainic Acid (alpha Kainic Acid, Cayman Chemicals, 78050) to increase neuronal activity. mCherry, mEmerald and mTagBFP2 emissions were sequentially detected with MBS-405, MBS 488/561/639 and BP 420-480 + BP 570-630, BP 420-480 + BP 495-550 and SBS SP 505 respectively. Z-stacks encompassing entire dendrites of 44.9 x 44.9 μm (1272 x 1272 pixels) were acquired.

Neurons expressing HaloTag-Sec61β/HaloTag-JPH3/HaloTag-STIM1/HaloTag-STIM2 (labeled with JF646 or JF635) with mCherry-JPH3/mScarlet-STIM2/mScarlet-CAAX2 or immunostained with AlexaFluor^TM^594, mEmerald-ER/mEmerald-Sec61β/mEmerald-Ca_V_2.1/mEmerald-Ca_V_2.2 or immunostained with AlexaFluor^TM^488 and mTagBFP2-CAAX2 for overlap analysis or contact sites interval analysis were performed in neurons at 18-22 DIV. Dendrites were identified by the presence of spines visualized with an appropriate plasma membrane marker. JF646, mCherry or mScarlet or AlexaFluor^TM^594, mEmerald or AlexaFluor^TM^488 and mTagBFP2 emissions were sequentially detected with MBS-405, MBS 488/561/639 and SBS LP 570, BP 420-480 + BP 570-630, BP 420-480 + BP 495-550 and SBS SP 505 respectively. Z-stacks encompassing entire dendrites of 44.9 x 44.9 μm (1272 x 1272 pixels) were acquired.

Time-series of neurons expressing mEmerald-STIM1 and HaloTag-Sec61β (+JF585) were acquired by detecting mEmerald and JF585 with MBS-405, MBS 488/561/639 SBS. 10-minute time-lapse videos of 67.3 x 67.3 μm (1584 x 1584 pixels) were acquired every 4 seconds. Tyrode buffer containing 1 μM thapsigargin (Sigma Aldrich, T9033) was perfused after 1 minute.

All Z-stacks were acquired using a piezo-motor and Nyquist-optimized z-step sizes (0.133 μm). Live imaging was performed at 37 °C and 5% CO2. Images were acquired and processed using automatic 2D or 3D Airyscan processing in ZEN software (Zeiss).

### Contact sites interval analysis

Contact sites interval analysis was conducted using intensity plot profiles of endogenous JPH3 tagged with mEmerald, HaloTag-JPH3, or mCherry-STIM2. These profiles were mapped along two-dimensional line scans tracing the dendritic plasma membrane from Airyscan z-stacks or 3D-lattice SIM volumes. Line traces generated from Fiji (Schindelin et al., 2012)^9^ were utilized to analyze contact site intervals. A Julia^10^ (version 1.8.5) Pluto.jl notebook (version 0.19.22), **imagej_peaks_and_intervals_with_area.jl**, was used to examine peaks in the raw intensity signals. The height, interval width, and average height of peak intervals were analyzed using data from CSV files loaded through CSV.jl. Peaks.jl was employed to find local maxima, minima, and peaks with a minimum prominence of 20 for endogenous tagged JPH3 and 200 for HaloTag-JPH3 and HaloTag-STIM2. The two local minima nearest each local maximum were used to create intervals around each local maximum. If local maxima were not identified as peaks due to the minimum prominence thresholds, the interval for each maximum was merged with the interval of the peak closest to the local maximum. The width of each interval was determined from the merged intervals around each peak. The area within each peak interval was numerically integrated to estimate the area under the curve, which could be assigned to each peak. To calculate the average height per peak, the area was divided by the interval width. Data were organized into a data frame using DataFrames.jl, containing columns such as peaks (peak location), value (peak height), prominence (prominence from Peaks.jl), area (integrated area within each interval), full_width (merged interval width around each peak), average_height (interval area divided by interval width), and peak intervals (distance between adjacent peaks). The data frames were then exported as Excel sheets using XLSX.jl. The Julia notebook, including the internal project specification and manifest can be found at the following Github link: https://github.com/JaneliaSciComp/Benedetti2024. The analysis used Julia version 1.9.3 was used with the following registered Julia packages: CSV.jl (v0.10.9), DataFrames.jl (v1.5.0), Peaks.jl (v0.4.3), Plots.jl (v1.38.8), PlutoUI.jl (v0.7.50), StatsPlots.jl (v0.15.4), XLSX.jl (v0.9.0), and Pluto.jl (v0.19.27). Statistical analysis was performed using the software GraphPad Prism (version 10.2.3).

### Contact sites periodicity analysis (periodicity.jl)

Local maxima (peaks) were detected using Peaks.jl with a minimum peak prominence set to 5% of the maximum fluorescence intensity profile from the Fiji line profiles exported to CSV. A synthetic signal was created by summing Gaussian profiles with a standard deviation of 71 nm (two pixels) centered at the locations of each detected peak. Fourier analysis was performed on this signal using FFTW.jl to identify the peak frequency in the Fourier power spectrum. The synthetic signal was then cross-correlated with a periodic Gaussian signal that had the same peak frequency. To determine the 95% confidence interval for the cross-correlation analysis, correlation with a white noise process was used as a model under the null hypothesis. The degrees of freedom of the signal, N, were estimated as the length of the signal in μm divided by the resolution of the image, ∼122 nm. Consequently, the 95% confidence intervals were set at 1.96/sqrt(N) and - 1.96/sqrt(N)^11^. The Julia notebook can be found at the following Github link: https://github.com/JaneliaSciComp/Benedetti2024.

### Raincloud Plots (raincloud.jl)

The package Pluto.jl was used via Julia to create an interactive notebook that generated raincloud plots. The interactive notebook loaded data from Excel .xlsx files at selected cell ranges. The legend names, title, x label, y label, number of plots, plot width, and plot orientation were customizable via a user interface. A Plots.jl recipe using the GR framework backend was used to draw a raincloud plot consisting of 1) a violin plot, 2) a box plot, and a 3) dot plot. The resulting plots were exported to PDF files. The packages Pluto.jl, 0.19.22, Plots.jl v1.38.8, Peaks.jl v0.4.3, XLSX.jl v0.9.0, PlutoUI.jl v0.7.50, StatsPlots.jl v0.15.4, HypertextLiteral.jl v0.9.4, and Measures.jl v0.3.2 were used to generate the raincloud plots. The Julia notebook can be found at the following Github link: https://github.com/JaneliaSciComp/Benedetti2024.

### Analysis of contact sites proximity to dendritic spines

Contact sites proximity to dendritic spines’ neck was analyzed using high-resolution 3D Lattice-SIM volumes of whole dendrites in neurons expressing HaloTag-JPH3, mScarlet-CAAX2, and mEmerald-Sec61β at 18-21 DIV. Once a focal plane was identified where the spine head, neck, dendritic shaft, laddered ER, and JPH3 puncta were all clearly discernible, line profiles were drawn from the spine neck to the closest contact sites along the dendritic plasma membrane using Fiji. Statistical analysis was conducted using GraphPad Prism.

### Analysis of the size, density, and overlap of ER-PM junction components

The size and number of hotspots for JPH3, Ca_V_2.1, Ca_V_2.2, and pCaMKII were determined using Fiji software (Schindelin et al., 2012)^9^. Each channel’s individual planes were processed with Gaussian Blur and background subtraction. A threshold was set after subtraction to identify and count objects. The number of objects was normalized to the surface of the dendritic profile by tracing its profile with the help of the plasma membrane marker to calculate hotspots/clusters density.

Object-based colocalization analysis was conducted using the DiAna plugin in Fiji (Gilles et al., 2017)^12^. In this analysis, the selected planes of the two channels of interest were filtered with Gaussian blur and then globally thresholded based on intensity. This allowed the plugin to identify the local maxima of each object as its center. The plugin then performed a center-to-center distance analysis and a contact surface analysis. Two objects were considered colocalizing when their center-to-center distance was less than 150 nm or when their overlap was greater than 50%. Statistical analysis was performed using the software GraphPad Prism.

### Volume electron microscopy and intracellular organelles segmentation

The FIB-SEM dataset covering the CA1 region of a 2-month-old C57/BL6 mouse was kindly provided by the Hess and Clapham labs. Round samples of the hippocampus were taken from 100 mm coronal sections of the brain obtained from a transcardially perfused mouse. The samples were prepared using a hybrid protocol as described in Sheu et al., 2022^13^. The FIB-SEM dataset comprising a portion of a 5-day-old female *Drosophila melanogaster* central brain (referred to as hemibrain) was described in Scheffer et al., 2020^14^. It can be openly accessible through the browser interface NeuPrint (https://neuprint.janelia.org). We utilized this dataset to examine the organization of intracellular organelles in invertebrate neurons and selected a neurite region of Mushroom Body Output neuron 1 (MBON1) for segmentation. Both samples were imaged by a customized Zeiss Merlin FIB-SEM system (previously described by Xu, C. S. et al. 2017^15^) and generated volumes composed of isometric 8 x 8 x 8 nm voxel sizes. For more information on the sample preparation, generation of volumet EM dataset, and post-processing, please refer to Sheu et al.^13^ and Scheffer et al.^14^. Both datasets were analyzed to identify intracellular organelles based on established morphological features. The organelles were then manually segmented using the 3D visualization and processing software platforms Amira-Avizo 6.7 (ThermoFisher) as described in Heinrich et al. 2021^16^ and Park et al. 2023^17^. The 3D segmentations of 3 mouse hippocampus medial dendrites were inspected for the organization of the ER, ER-PM junctions, and mitochondria.

Quantification of consecutive ER-PM contact site intervals was performed by measuring the distance between consecutive contacts using ER tubules as a guide with Amira-Avizo. The 3D segmentation of *Drosophila* neurite was inspected for the organization of the ER, ER-PM junctions, microtubules, and mitochondria. ER-PM contacts were identified as voxels where the distance between the ER and PM was less than 30 nm.

### Hypotonic cell swelling

Primary rat hippocampal neurons were infected at 5 DIV to co-express HaloTag-Sec61β (labeled with JF646 HaloTag ligand), mCherry-JPH3, and mTagBFP2-CAAX2 to identify ER membranes, ER-PM junctions, and the plasma membrane, respectively. Neurons at 21 DIV were imaged with Airyscan microscopy prior to hypotonic treatment to confirm the presence of a fine, tubular ER network extending throughout dendrites, with regular mCherry-JPH3 contacts. Dendrites were identified by visualizing dendritic spines using the PM marker mTagBFP2-CAAX2. Neurons were then incubated with hypotonic swelling media (20% Tyrode buffer in water yielding a ∼63 mOsM solution) at 37°C for 10 minutes. The ER’s fine tubular network transformed into numerous micrometer-sized large intracellular vesicles composed of mEmerald-Sec61β-positive membranes (as described in King et al., 2020^18^) that remained regularly distributed along the imaged dendrite. A 405 nm laser was used to excite mTagBFP2-CAAX2, a 561 nm laser for mCherry, and a 639 nm laser for JF646 HaloTag-ligand. Fluorescence emission was detected with appropriate bandpass filters MBS-405 and MBS488/561/639 + SBS LP 560 nm, BP 420-480 + 570-620 nm + SBS LP 560, respectively. Fields of view of 67.3 x 67.3 µm (1908 x 1908 pixels) at 0.13 µm z-steps were acquired. The interval of mCherry-JPH3 contact sites upon incubation with hypotonic swelling media was performed as described previously, mapping intensity profiles of mCherry-JPH3 along two-dimensional line scans tracing the dendritic plasma membrane and analyzed with a Julia Notebook for contact interval analysis and the software GraphPad Prism.

### Immunofluorescence staining of primary neurons

Primary rat neurons were fixed in PBS (Fisher Scientific, 70-011-069) with 4% paraformaldehyde (Electron Microscopy Sciences, 50-980-487), 0.1% glutaraldehyde (Electron Microscopy Sciences, 16019), and 4% sucrose (Fisher Scientific, S5-500) for 30 minutes at 37°C. Subsequently, cells were washed three times with PBS 1x and incubated with 0.01% sodium borohydride (Sigma Aldrich, 213462) for 7 minutes. After three washes with PBS (Fisher Scientific, 10010023), neurons expressing a protein of interest fused to HaloTag were incubated with JF dyes at 100 nM for 30 minutes in PBS. After three rinses with PBS, the cells were incubated with a 0.2% Triton-X100 solution (Sigma Aldrich, 93443) in PBS for 10 minutes. To prevent non-specific binding, a blocking step was performed for 30 minutes with BlockAid™ Blocking Solution (ThermoFisher, B10710). The primary antibodies were then added in BlockAid solution, and the cells were incubated overnight at 4°C. The following primary antibodies were used: rabbit polyclonal anti-Ca^2+^ channel P/Q-type alpha-1A antibody (1:400 dilution, Synaptic Systems 152203); rabbit monoclonal anti-phospho-CaMKII (Thr286) (D21E4) (1:800 dilution, Cell Signaling Technology 12716); mouse monoclonal anti-CaMKII alpha (1:800 dilution, Thermofisher, 6G9, MA1-048); rabbit polyclonal anti-SERCA2 ATPase (1:400 dilution, Abcam, ab3625); mouse monoclonal anti-Ryanodine Receptor 34C (1:200 dilution, Abcam, ab2868). After three rinses with 0.1% Triton in PBS for 10 minutes, the cells were incubated with the appropriate secondary antibody in BlockAid solution for 2 h. The following secondary antibodies were used: Goat anti-Mouse IgG (H+L), Superclonal™ Recombinant Secondary Antibody, Alexa Fluor 488, Thermo Fischer, A28175; Donkey anti-Rabbit Alexa Fluor 488, Jackson Laboratory, 711-547-003; Donkey anti-Mouse Alexa Fluor 594, Jackson Laboratory, 715-587-003; Donkey anti-Rabbit Alexa Fluor 594, Jackson Laboratory, 711-587-003. Finally, the cells were rinsed three times with 0.1% Triton in PBS for 5 minutes and three times with PBS for 5 minutes before being imaged in PBS. The cells were then mounted in ProLong™ Diamond Antifade Mountant (ThermoFisher, P36961) for long-term storage.

### JPH3 knockdown and validation

Primary rat hippocampal neurons at DIV3 were infected with pLKO.1-TRC mTagBFP2 (Addgene, plasmid #191566) to express JPH3 shRNA (from OrigGene, TL705996) and the same lentiviral construct encoding a scramble shRNA was used as a control. After infection neurons at 21 DIV were imaged, collected for verification by immunoblot, or FAC sorted for verification by RT-PCR.

For immunoblot experiments, primary neurons were homogenized in RIPA buffer (Cell Signaling Technologies, Cat # 9805S) with cOmplete™, Mini, protease Inhibitor Cocktail (Roche, 11836153001) and 100 mM PMSF (Cell Signaling Technologies, Cat # 8553S) on ice. After lysis, samples were centrifuged at 265,000g at 4 °C for 1 hour and the supernatant was collected. Total protein in the lysates was determined by Pierce™ BCA Protein Assay Kit, (ThermoFisher, 23227). Samples were then mixed with Laemmli buffer (Millipore Sigma, S3401) and resolved by SDS-Polyacrylamide Gel Electrophoresis (SDS-PAGE) on NuPAGE™ 4 to 12%, Bis-Tris, 1.0 mm, Mini Protein Gel, 10-well (ThemoFisher, NP0321BOX) and transferred to polyvinylidene fluoride (PVDF) membranes (Biorad, 1704156EDU). PVDF membranes were blocked with 5% nonfat dried milk in Tris-buffered saline (TBS) pH 7.4 with 0.1% Tween-20 (Sigma Aldrich, P9416) and probed with a polyclonal rabbit anti-JPH3 antibody (1:1000 dilution, Abcam, ab79063), and rabbit polyclonal anti-GAPDH antibody (1:5000 dilution, Abcam, ab9485). Primary antibody binding was detected with horseradish peroxidase-conjugated secondary antibodies (Goat Anti-Rabbit IgG H&L (HRP), Abcam, ab205718) and visualized by chemiluminescence (Thermo Fisher Scientific, SuperSignal West Pico PLUS, 34578). The blots were analyzed using ImageJ>Analyze>Gels. To calculate the percent knockdown, normalization factors were used to measure the immunoblots. The results are displayed in the representative images as ’relative total protein’. These percent knockdown calculations are based on the outcomes of at least three independent experiments.

For RT-PCR, RNA extraction was performed on 3 independent experiments using an adaptation of the protocol created by Dr. Peter White of the Research Institute at Nationwide Children’s Hospital (NCHRI), which utilizes the RNeasey® Mini Kit (Qiagen Cat# 74106). Total RNA was quantified via NanoDrop Onec (Thermo Scientific), and the extraction products were adjusted to equivalent total RNA concentrations. cDNA first strand synthesis was performed using the SuperScript™ IV VILO™ Master Mix (Thermo Fisher Scientific Cat# 11756050) reverse transcriptase kit. In brief, the qRT-PCR reaction mix was prepared by combining the reverse transcription products with either endogenous control or Jph3 primers (rGAPDH: Forward CAACTCCCTCAAGATTGTCAGCAA and Reverse GGCATGGACTGTGGTCATGA; Jph3: Forward AAGTCCAGCACGGGCTCAG and Reverse TTGCTTCCGTGGTCACAACA) and the PowerTrack™ SYBR™ Green Master Mix (With the ROX passive reference dye). qRT-PCR was then performed using a QuantStudio 7 Pro (Applied Biosystems) in standard mode. A melt curve was generated to confirm uniform amplification. Using the Applied Biosystem’s Design and Analysis (DA2) software suite, the relative expression of Jph3 was calculated with the ΔΔCT method. Statistical significance was determined by unpaired t-test with the software GraphPad Prism.

### Detection of pCaMKII and CaMKIIα

Neuronal stimulation and detection of pCaMKII or CaMKIIα were performed using the protocol developed by the Tsien lab^19^. To visualize the plasma membrane of dendrites and spines, mTagBFP2-CAAX2 was expressed in neurons together with mCherry-JPH3 to map the position of contact sites along dendrites. At 21DIV, neurons were equilibrated in Tyrode solution (Tyrode 5 mM K^+^: 150 mM NaCl, 5 mM KCl, 2 mM CaCl_2_, 2 mM MgCl_2_, 10 mM HEPES and 10 mM glucose, pH 7.4) containing 0.5 μM TTX (Tetrodotoxin citrate, Abcam ab120055) to block action potentials, 10 μM CNQX (Sigma Aldrich, C239) to block AMPA receptors, and 10 μM AP5 (Sigma Aldrich, A5282) to block NMDA receptors for 15 minutes at 37°C and 5% CO_2_. Neurons were stimulated using 90 mM K^+^ Tyrode solution (65 mM NaCl, 90 mM KCl, 2 mM CaCl_2_, 2 mM MgCl_2_, 10 mM HEPES and 10 mM glucose, pH 7.4). The solution was supplemented with 10 μM Nimodipine (Abcam, ab120138), 500 nM omega-Agatoxin IVA (Abcam, ab120210), and 2 μM ω-conotoxin GVIA (Tocris, 1085). These were added before, during the stimulation and recovery process to inhibit voltage-dependent calcium channels (Ca_V_ Block). After a 45-second recovery in 5 mM Tyrode, neurons were fixed and immunolabelled using a rabbit monoclonal anti-phospho-CaMKII (Thr286) (D21E4, Cell Signaling) or a mouse monoclonal anti-CaMKIIα (6G9, Thermofisher, MA1-048). This was done as described previously, with the use of Donkey anti-Rabbit AlexaFluor^TM^488 and Goat anti-Mouse AlexaFluor^TM^488 secondary antibodies. Neurons were imaged in PBS after fixation using Airyscan microscopy. Sequential detection of mCherry, AlexaFluor^TM^488, and mTagBFP2 emissions with MBS-405, MBS 488/561/639 and, BP 420-480 + BP 570-630, BP 420-480 + BP 495-550 and SBS SP 505 respectively was done. Z-stacks were taken at 0.13 µm z-steps, covering entire dendrites or cell bodies of 44.9 x 44.9 μm (1272 x 1272 pixels). The identification of JPH3 contacts and pCaMKII hotspots was performed using Fiji software. Individual planes of each channel were processed with Gaussian Blur and background subtracted. For pCaMKII signal, the background was measured in a region of interest (ROI) located at the center of the dendrite. After subtraction, a threshold was set to identify and count objects. The number of objects was normalized for the surface of the dendritic profile measured by tracing its profile with the help of the plasma membrane marker. Colocalization analysis was performed with the DiAna plugin in Fiji. Statistical analysis was performed using the software GraphPad Prism.

### Two-photon glutamate uncaging

Sprague-Dawley rats (postnatal P0) were obtained from the in-house animal core facility of the Max Planck Florida Institute for Neuroscience. Hippocampal regions were dissected and dissociated with the Worthington Papain Dissociation Kit (Worthington Biochemical Corporation, stored at 4 °C) with a modified manufacturer’s protocol. Briefly, the digestion is done once for 30 min. Following trituration, the cells are centrifuged at 300 rcf for 5 minutes, resuspended in 2.4 ml of resuspension buffer (1.44 ml EBSS, 160 μl albumin-ovomucoid inhibitor, and 80 μl DNase), and gently added to the top of 3.2 ml albumin-ovomucoid inhibitor to form a gradient. The cells were then forced through the gradient by centrifugation at 600 rpm for 12 minutes. Neurons were plated on poly-D-lysine coated coverslips mounted on Mattek dishes at a density of 39,000-52,000 cells/cm^2^. Cultures were maintained in NGM (Neurobasal-A Medium, 2% B-27, and 1% Glutamax, from Gibco, ThermoFisher Scientific, sterile filtered using a 0.1 μm filter and stored at 4 °C) at 37°C and 5% CO_2_. Cells are fed with 1 ml NGM 3 - 4 h after plating and with 0.5 ml NGM 3 days after plating. After that, feeding is performed every 3 - 4 days by removing 0.3 ml conditional medium and adding 0.5 ml NGM. Transfections were performed 12-15 days after plating by magnetofection using Combimag (OZ biosciences) and Lipofectamine 2000 (Life Technologies) according to the manufacturer’s instructions. Unless specified otherwise, all reagents were purchased from Sigma, and all stock solutions were stored at -20°C.

Live cell imaging was conducted 13-20 days after neuronal cell culture plating. All experiments were performed in a modified E4 imaging buffer containing: 120 mM NaCl, 3 mM KCl, 10 mM HEPES (buffered to pH 7.4), 4 mM CaCl_2_, and 10 mM Glucose, lacking MgCl_2_. Imaging was performed using a custom-built inverted spinning disk confocal microscope (3i imaging systems; model CSU-W1) attached to an Andor iXon Life 888. Image acquisition was controlled by SlideBook 6 software. Images were acquired with a Plan-Apochromat 63x/1.4 NA. Oil objective, M27 with DIC III prism, using a CSU-W1 Dichroic for 488/561 nm excitation with Quad emitter and individual emitters, at laser powers 2.00 mW (488 nm) and 2.65 mW (561 nm) for spine stimulation measurements. During imaging, the temperature was maintained at 37 °C using an Okolab stage top incubator with temperature control.

For ER calcium spatial profile measurements, neurons were transfected with ER-GCaMP6-210 and PSD95-mCherry plasmid constructs. PSD95-mCherry fluorescence was used to identify spines for stimulation by two-photon glutamate uncaging. Before glutamate uncaging, neurons were treated with 1 μM TTX (Citrate salt, 2 mM stock made in water) and 2 mM 4-Methoxy-7-nitroindolinyl-caged-L-glutamate (MNI caged glutamate) (Tocris Bioscience, 100 mM stock made in modified E4 buffer) in modified E4 buffer lacking Mg^2+^ (see above). Glutamate uncaging was performed using a multiphoton-laser 720 nm (Mai TAI HP) and a Pockel cell (ConOptics) for controlling the uncaging pulses. Spines that were at least 70 μm away from the cell body were chosen for uncaging experiments. To compare different ER and cytosolic calcium spatial profiles in dendrites, uncaging protocols of 6 uncaging pulses at 0.2 Hz with 100 ms pulse duration per pixel at 7.2 mW power were used. For Ryanodine treatment experiments, neurons were treated with 100 μM Ryanodine (Tocris Bioscience, 25 mM stock made in DMSO) for 5 min, and imaging continued as described above. For cytosolic Ca^2+^ spatial profile measurements, neurons were transfected with the cytosolic Ca^2+^ sensor RCaMP1.07. Spines were stimulated by 6 uncaging pulses at 0.2 Hz with 100 ms pulse duration per pixel at 7.2 mW laser power to compare with ER calcium spatial profile. Image analysis was done using Fiji. First, images were averaged across the 6 uncaging time points unless specified otherwise. Then, a line of 1 μm thickness flanking up to 40 μm equidistant on either side of the stimulated spine was drawn manually using the ‘segmented line’ tool along the dendrite. The stimulated spine position was marked as 0 μm. The negative μm distances to the left of the stimulated spines denote the direction of the cell body, whereas the positive μm distances to the right of the stimulated spines denote the direction of the dendritic tip. The fluorescent profile of the line drawn along each dendrite was measured by the ‘plot profile’ function to obtain the spatial profile. ER-GCaMP6-210 channels were used to measure the ER Ca^2+^ spatial profile, and the RCaMP1.07 channels were used to measure the cytosolic Ca^2+^ profile. The peak detection analysis of ER-Ca^2+^ release profiles was performed, mirroring the original signal to allow for periodic analysis. Type I discrete cosine transform via FFTW was performed. The initial peak analysis is done by analyzing the high-pass filtered discrete signal. This was done using the Peaks.jl analysis from Allen Hill (https://github.com/halleysfifthinc/Peaks.jl). Selection of peaks is then possible based on the value of the high pass filtered discrete signal. This analysis was performed with the Julia Pluto.ij notebook**: fft_analysis.jl** that can be found on Github: https://github.com/JaneliaSciComp/Benedetti2024.

### Resource Availability

#### Lead Contact

Requests for further information and reagents should be directed to and will be fulfilled by the lead contacts at.

#### Material Availability

Plasmids generated in this study will be available in the Addgene repository.

#### Data and Code Availability

Julia Pluto.jl notebooks are available on Github at the following link: https://github.com/JaneliaSciComp/Benedetti2024.

## Notes

### Competing Interest Statement

The authors have declared no competing interest.

